# Multi-Stage Graph Attention Networks for Interpretable Alzheimer’s Disease Classification from Genome-Wide Association Data

**DOI:** 10.64898/2026.04.06.716790

**Authors:** Ankita Saxena, Christopher Gaiteri, Stephen V Faraone

**Affiliations:** Department of Neuroscience and Physiology, State University of New York – Norton College of Medicine at Upstate Medical University, Syracuse, New York, USA; Department of Psychiatry and Behavioral Sciences, State University of New York – Norton College of Medicine at Upstate Medical University, Syracuse, New York, USA; College of Medicine, MD program, State University of New York – Upstate Medical University, Syracuse, New York

## Abstract

**Background:** Genome-wide association studies have identified numerous variants associated with neuropsychiatric disorders. Although some significant loci can carry substantial risk, as in Alzheimer’s Disease, the remaining genetic variance is distributed across many small-effect loci. Polygenic risk scores (PRS) aggregate this risk but do not capture epistatic interactions, and offer limited biological interpretability and predictive accuracy. Computing gene level risk scores and integrating known or statistically validated gene-gene associations has the potential to increase interpretability and/or accuracy. Graph Neural Networks (GNNs) can leverage graph structured genetic data that models potential epistatic interactions to achieve these goals.

**Methods:** We developed a three-stage Graph Attention Network (GAT) classifier using individual-level GWAS data from 7,358 participants across seven Alzheimer’s Disease Center cohorts. Nodes were defined as genes, with risk scores from AD and 11 genetically correlated phenotypes serving as features. We evaluated two graph construction strategies: gene co-expression networks derived from hippocampal transcriptomic data and curated pathway-based graphs. Additionally, a bilinear context module was incorporated to capture global gene-gene interactions beyond the graph topology. In Stage 1, a GNN encoder was trained on the graphs; Stage 2 injected PRS for non-coding SNPs after the encoder to better capture genetic risk via transfer learning, and Stage 3 applied adversarial training with gradient reversal for ancestry debiasing. GNN predictions were ensembled with whole-genome PRS using elastic net regression.

**Results:** The best-performing GNN model — a GAT with bilinear context operating on the pathway graph — achieved an AUROC of 0.78 (95% CI: 0.75–0.80). Ensemble models combining Stage 2 or 3 GNN logits with whole-genome PRS achieved an AUROC of 0.82 (0.79–0.84), outperforming PRS alone (0.80). GxI attribution and additional explainability analyses revealed stage-specific biological signals, some of which re-capitulated known gene-phenotype associations and others which may reflect potential new areas of inquiry.

**Conclusion:** A multi-stage GAT framework captures complementary, non-additive genetic signal that, when ensembled with PRS, improves the accuracy of AD classification. Post-hoc explainability analyses yield biologically interpretable gene networks, supporting the utility of graph-based deep learning for dissecting complex genetic architectures.

## Introduction

Uncovering the contributions of genetics to the development of diseased phenotypes has been a longstanding objective in biomedical research. Different tools have been used in this quest, including Genome Wide Association Studies (GWAS), which, since their advent in 2005, have enabled scientists to discover associations between genetic variants and numerous traits (Ikegawa 2012; Uffelmann et al. 2021). These efforts have shown that while some diseases are mendelian, many others, including psychiatric disorders, diseases of metabolism, and inflammatory pathologies qualify as complex genetic traits (Ali 2013; Cleynen et al. 2016; Dedmon 2020; Sullivan et al. 2012; Sullivan and Geschwind 2019). Critically, the expanding size of GWAS as well as improved typing of genetic variants over the last decade has facilitated the discovery of many more small nucleotide polymorphisms (SNPs) that are linked to complex disorders; however, these individual variants are still insufficient to explain the phenotypic variance, or heritability, across the population (Demontis et al. 2023; Pettersson et al. 2019). In contrast, methods that incorporate many GWAS SNPs have been more successful; however, a gap remains between the heritability estimates obtained from these multi-SNP methods and the heritability values found vis a vis twin studies (de Los Campos et al. 2015; Young 2019). Altogether, these findings suggest complex genetic architectures that include polygenicity and genetic heterogeneity as well as potential roles for epistasis and gene-environment interplay (Crouch and Bodmer 2020; Wendt et al. 2020).

Among the best known of the multiple-SNP/“whole genome” approaches are polygenic risk scores (PRS), which sum up allelic risk at numerous loci throughout the genome to create single scores that quantify genetic risk (Choi et al. 2020; Plomin and von Stumm 2022). PRS have shown some efficacy as predictors in research studies and are being tested for potential clinical use (Hao et al. 2022). Variations on PRS, including efforts that have expanded PRS use to include genetically correlated disorders, or scores for gene sets found to be associated with a trait of interest, have further enhanced predictive accuracy (Barnett et al. 2022). Nonetheless, areas for improvement remain; as single values, PRS cannot provide insight into which specific loci are major contributors of risk for a person or even a population; such information could help elucidate pathologic mechanisms (Plomin and von Stumm 2022). In tandem, PRS are a linear sum of allelic risk and cannot factor in possible interactions among genetic loci.

Consequently, more granular approaches that account for epistasis may offer improvements in predictive accuracy and/or further our understanding of disorder etiology (Phillips 2008). At the SNP level, epistasis can be challenging to detect or incorporate; exhaustive testing for interactions between sets of SNPs ranges from computationally expensive to infeasible depending on the set size, while interactions between loci with high effect allele frequency are likely to be correlated with the loci genotypes themselves and therefore, folded into narrow sense additive heritability (Balvert et al. 2024; Hemani et al. 2013; Niel et al. 2015; Russ et al. 2022). However, prior research has shown that networks of interactions between biological molecules are ubiquitous and essential for cellular and higher order function (Barabási and Oltvai 2004; Ma and Gao 2012). One approach to incorporating this framework is to adopt data representations that enable the evaluation of candidate epistatic interactions at the level of functional genetic units, such as genes. In parallel, existing knowledge such as statistical correlations of gene expression or pre-existing pathways in relevant tissues can be used to provide biological context. These sources of information offer both local insight into relationships among individual genes and broader context regarding groups of genes that are known to interact, thereby guiding the selection of interactions to be tested.

Deep learning neural networks are an alternative approach to conventional statistical methods that dynamically learn associations between input features to create new representations useful for making predictions (LeCun et al. 2015). Graph Neural Networks (GNNs) are specialized for graph structured data; convolutions are performed on the graphs, allowing the generation of graph level embeddings that reflect context sensitive aggregations of input features for use in downstream tasks (Bruna et al. 2013; Kipf and Welling 2016; Saxena et al. 2025). Graph Attention Networks (GATs) are a unique type of GNN in which attention scores are computed for each edge that connects any two nodes, reflecting the relative importance of the association to the updated node representation (Veličković et al. 2017). This allows the model to dynamically weight the importance of different edges when generating new representations of the inputs. However, GATs are limited in their ability to compute attention and therefore identify useful relationships, to the nodes that are connected by edges. Although multiple graph layers allow for an individual node to incorporate information from neighbourhoods up to *k* nodes away, this can result in over smoothing, i.e. node representations becoming very similar, which provides a practical constraint. Consequently, global graph representations can also be used to capture pairwise interactions between nodes that are not connected. Post-hoc explainability analysis of a trained GAT classifier, via gradients or even attention scores, can be used to extract important input features, gene-gene associations, and ultimately define biological networks useful in the prediction task (Yuan et al. 2022).

The objective of this work is to develop a GAT based classifier that operates on graph structured GWAS data and uses post-hoc methods to evaluate whether the incorporation of epistatic interactions and broader biological context assists in distinguishing between cases and controls. We construct graphs such that each node represents a gene, and each edge reflects a gene-gene interaction; we experiment with two methods of injecting context via graph construction. These use (1) gene co-expression networks derived from transcriptomic data, and (2) a set of previously identified pathways and relationships that are pruned/modified to optimize graph geometry. We also assess the value of including additional, gene risk scores derived from summary statistics for genetically correlated disorders, as determined by cross-trait Linkage Disequilibrium (LD) Score Regression, as additional node features in our predictive models (Bulik-Sullivan et al. 2015).

## Methods

### Data Acquisition and Pre-Processing

Individual level genotype and phenotype GWAS data were obtained from the seven sets of Alzheimer’s Disease Center (ADC) genotyped subjects used in a previous GWAS analyses (Kunkle et al. 2019). Summary statistics from Wightman *et al* that excluded the 23andme and UKBiobank samples, and also did not use data included in our analysis were used for posterior effect size adjustment via PRS-CS and for finding genetically correlated disorders via LD score regression through the Complex-Traits Virtual Genetics Lab (CTVGL) interface (Cuéllar-Partida et al. 2019; Ge et al. 2019; Wightman et al. 2021). To appropriately label the summary statistics with SNP IDs prior to genetic correlation analysis, the data was harmonized with a GRCh37 FASTA reference file, a VCF was created, and BCFTools were deployed to map SNPs onto dbSNP and ultimately collapse them to one SNP per chromosome/base-pair combination (Auton et al. 2015; Danecek et al. 2021).

Individual genotype data and all summary statistics used with PRS-CS were harmonized with the 1000 Genomes Phase 3 data on the GRCh37 build (Auton et al. 2015). Imputation and quality control steps were followed as previously described, with filtering being done for MAF < 1%, genotype missingness >5% and a Hardy-Weinberg equilibrium p-value <1e-06 using Plink v2.0 (Chang et al. 2015; Hou et al. 2022). The Average Age of Onset/Average Age of Examination (AAO/AAE) covariate was scaled and used to generate quantiles; the R caret package was used to create stratified groups based on AAO/AAE, sex, and phenotype (Hou et al. 2022; Kuhn 2008). 70% of samples were assigned to the training and 15% each to the test and validation subsets, resulting in respective dataset sizes of 5152 individuals for the training set, 1101 individuals for test set and 1105 individuals for validation. Principal components analysis was run on the training set via the “PCA approx” command on Plink v2.0 and the top 10 principal components (PCs) identified; the loadings from the training set were then used to compute the PCs for the test and validation sets.

Eleven genetically correlated phenotypes with h^2^ > 0.1, n > 10000, and absolute RG value of >0.10 were identified using the CTG-VL’s LD score regression analysis feature with formatted Wightman *et al* summary statistics inserted as input: Caudate Volume, Pallidum Volume, Thalamus Volume, Putamen Volume, Red Blood Cell (Erythrocyte) Distribution Width, Apolioprotein A, HDL Cholesterol, Started Insulin Within a year of Diabetes Diagnosis, Schizophrenia, Albumin, and Fluid Intelligence Score(2014; Hibar et al. 2015; Howrigan et al. 2023). As UK Biobank data had been excluded from the AD summary statistics, all publicly available summary statistics were downloaded and utilized for analysis. Subsequently, SNP effect sizes were adjusted for linkage disequilibrium using Bayesian posterior shrinkage via PRS-CS and a 1000 Genomes European LD reference panel; this approach was used in lieu of Plink’s “clump and threshold” approach to maximize retention of SNPs (Auton et al. 2015; Gaunt et al. 2007; Ge et al. 2019).

SNP to gene mapping was performed via a multi-level, sequential pipeline that removed mapped SNPs at each step. The pipeline first incorporated functional evidence to assign SNPs to genes; the remaining SNPs were then assigned using physical proximity, with each SNP in the analysis being mapped to only one gene. All SNPs that had posterior effect sizes inferred by PRS-CS and were available in the AD summary statistics were annotated using the Variant Effect Predictor (VEP) for functional consequences and gene symbols and types(McLaren et al. 2016). The list of sequence ontology terms used to define consequences included “stop_gained”, “stop_lost”, “frameshift_variant”, “splice_acceptor_variant”, “splice_donor_variant”, “missense_variant”, and “start_lost.” SNPs with any of these terms in the consequence field of the VEP output were assigned to the corresponding gene and removed from the overall SNP set. Gene annotations from GENCODE release 19 (GRCh37.p13) were subsequently used to create a BED file for of Transcriptional Start Sites (TSS); the bedtools window command was then applied to identify SNPs that fell within ± 2 kilobases of a TSS (Mudge et al. 2024) and map them to a gene. To help assign the remaining SNPs, an annotation file was created using the training set Plink v1.9 bfile, a GRCh37/hg19 based gene location file, and MAGMA software with an extended gene window of 35 kilobases upstream and 10 kilobases downstream from the gene (de Leeuw et al. 2015; Purcell et al. 2007; Zhang 2022). SNPs were mapped to the closest gene by proximity; SNPs that remained unmapped at this point were deemed intergenic SNPs. Gene level risk scores were computed for each phenotype using the SNPs assigned to each gene and the adjusted effect sizes via Plink 2.0’s scoring method (Figure 1a). PRS were also obtained for each phenotype using all SNPs from the PRS-CS outputs and Plink v2.0. To quantify genetic risk attributable to intergenic SNPs, SNPs used for genes included in each graph were summed up and subtracted from the genome wide set of SNPs; the remaining SNPs were used to construct a weighted score. These intergenic PRS were injected as a graph level feature into the GNN (Figure 1b).

**Figure 1:**
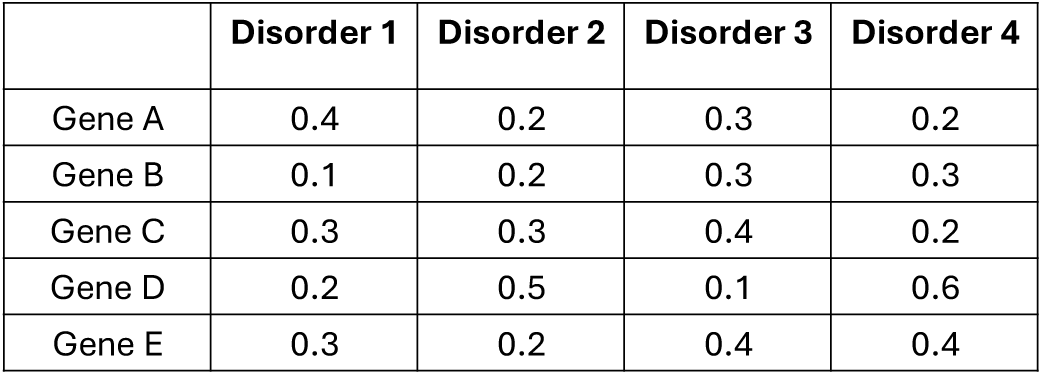
Model Architecture and Data Processing Pipeline for Pathway-Based AD Classification. Figure 1a: Node feature matrix: gene-level risk scores across AD and correlated disorders

**Figure 1b:**
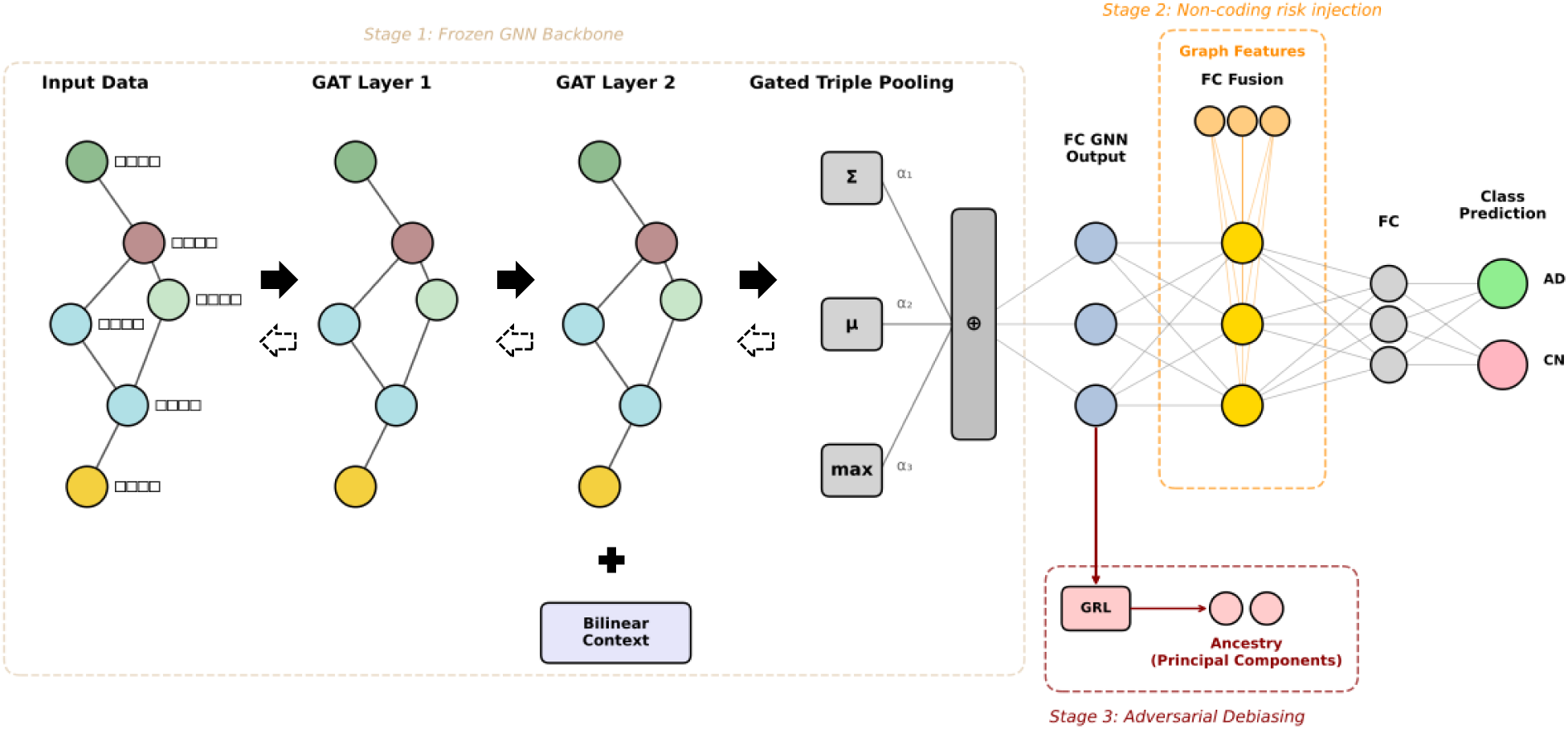
Multi-stage GNN architecture with adversarial ancestry debiasing

Adjacency matrices to define graph structures were obtained through several sources. Normalized aggregated gene expression data from post-mortem control and AD tissue from the hippocampus was obtained from a previous analysis (Hou et al. 2022). Pearson’s correlation coefficients were computed for the gene expression data using R version 4.4.2; values were rounded to the second decimal point and the absolute values taken. The correlation matrix was looped over to extract the top 0.125, 0.25, and 0.375% of correlations for each gene; all others were reduced to zero. Adjacency matrices were filtered to exclude genes that did not have scores for all phenotypes, ultimately resulting in the inclusion of 16, 366 genes for the gene expression graphs.

Pathway Graph generation began with the same 16,366 genes used in the gene expression graphs; genes were mapped to Entrez IDs using the biomaRt package, with an alias to canonical symbol dictionary being used to rescue initially unmapped genes (Durinck et al. 2009; Sayers et al. 2022). Genes that did not map despite the alias rescue were removed and their genomic risk was added to the intergenic risk sums; the final Pathway Graph thus included 16, 352 genes. Edges were then constructed via a tiered mechanism; for the first tier, regulatory/interaction edges from KEGG pathway graphs were extracted using the graphite package; second, genes co-participating in the same Reactome pathway (capped at 50 genes/reaction) were connected; third, pairwise edges for genes sharing more than 2 pathways were constructed. (Gillespie et al. 2022; Kanehisa et al. 2023; Sales et al. 2012). After the first three steps, still-isolated genes were subject to rescue mechanisms to connect them: in the fourth step, they were rescued if they shared at least 1 KEGG/Reactome pathway; next, remaining isolated genes were rescued using Gene Ontology Biological Processes terms (terms were limited to those encompassing 2 to 200 genes); finally, gene expression data from hippocampal Alzheimer’s Disease samples was used to connect remaining isolates with their top 5 correlated genes (2021). At this step, all genes were connected to at least one other gene, with a total of 3,733, 420 undirected edges and a mean node degree of 228 and maximum of 3521.

Given that dense graph connectivity can degrade GNN performance via over-smoothing, or the convergence of feature representations, as well as the propagation of noisy or irrelevant signals, we sought to prune the Pathway Graph prior to use (Chen et al. 2020; Li et al. 2018). To do this, we computed Forman-Ricci curvature for all edges in the initial graph and pruned approximately 75% of edges with curvature less than −206, while ensuring that each node had at least two edges (Forman 2003; Sreejith et al. 2016). This yielded a graph with approximately 936,742 edges, a mean and maximum node degree of 57.37 and 355 respectively, and a positive mean curvature of 22.17. However, this left approximately 12% of nodes having the minimum of 2 edges, which can lead to over-squashing and prevent the propagation of distant information. As a result, we also performed curvature guided rewiring inspired by discrete Ricci flow (Chen et al. 2020). At each iteration, we recomputed Forman-Ricci curvature on the pruned graph and sampled an edge with probability proportional to the negative curvature, thereby identifying structural bottlenecks. We then added shortcut edges between nodes that were 2 hops away, creating alternative pathways for the flow of information. This reduced over squashing by bypassing high traffic edges while increasing local clustering. We performed 20 iterations where 50 edges were added each time, increasing the total number of edges in the graph to 938, 672. We also generated a membership matrix for each node and edge sources and saved it for downstream analysis.

### GNN Architecture and Design

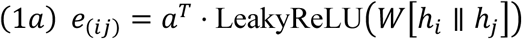

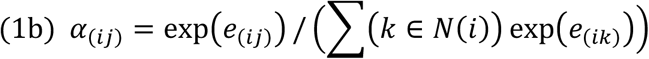

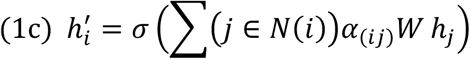

The GATConvV2 layer developed by Brody *et al* was used for its ability to compute attention for each edge on a node by node level vs graph level (Brody et al. 2021; Veličković et al. 2017). For each connected pair of genes in the graph, (*i, j),* we computed an attention coefficient (equation 1a) where **h***i* and **h***j* are concatenated feature representations of genes *i* and *j,* **W** is a learned weight matrix, and **a** is a learned attention vector. These coefficients were subsequently normalized (equation 1b), and can be interpreted as the learned importance of gene *j* in the context of gene *i*, before they are used to generate an updated representation for each (**h***i*′) by aggregating information from its neighbors in an attention weighted operation (equation 1c).

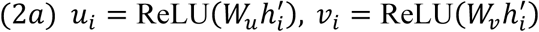

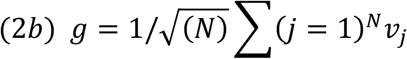

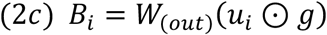

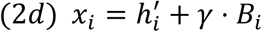

Although attention-based message passing can capture localized gene-gene interactions, global interactions that are not defined by the graph may also contribute to genetic risk. Subsequently following all graph layers, the updated gene representations were projected onto two complementary subspaces, wherein **ui** captured gene-specific features, and **vi**, a global context representation (equation 2a). The global context vector aggregates information across all genes in the graph, with scaling being applied to maintain stable gradients (equation 2b). Local and global features are then combined via element-wise multiplication, allowing the model to learn bilinear interactions (*B_i_*) between a gene’s local state and the broader genomic context (equation 2c). Finally, the contextual information is incorporated into the gene level feature representation (equation 2d) with *γ* being a learned parameter that scales the influence of the bilinear interactions. We tested models both with and without bilinear interactions.

The GNN model was trained in three stages (Figure 1b). The first stage of the model consisted of a GNN encoder that trains on the input graph with the number of convolutions, hidden dimensions, and other hyperparameters determined via Bayesian tuning. Following the completion of the GAT layers, sum pooling, mean pooling, and max pooling were done for each hidden dimension to capture different types of signal. These pooled values for each component were concatenated to form graph wide embeddings; to prevent sum pooling from dominating, values from each pooling output were scaled, with the scales treated as trainable parameters.

Graph wide embeddings were, in turn, fed into a dense fully connected layer (FCNN1), with maximum dimension of 128, and then, another tunable fully connected layer (FCNN2) which could alternatively serve as a bottleneck or a projector, depending on the hyperparameter value, before feeding into a final layer (FCNN3) that led to class status prediction via softmax. GAT convolution and linear layers were followed by ReLU activation functions; to prevent overfitting and maximize sparsity, dropout was also used. A binary cross entropy function was used for the loss function in training in conjunction with the Adam optimizer (Kingma and Ba 2014). Class weights were deployed to account for class imbalance.

To estimate how well the input graphs predicted global ancestry, a known confound in genetic modelling, we created a dual headed encoder model, where the outputs of the GNN encoding layer were also used to predict the top 10 PC projections. A stop gradient was placed after the FCNN1 that follows the graph embeddings to prevent the use of gradients from the ancestral head to update weights while still allowing the regression head to maximize performance. For this estimation approach, the tuned hyperparameters from stage 1 were used and the mean squared error (MSE) loss and PC correlations were recorded.

In stage 2 of the model, intergenic PRS were injected as graph level features after the second fully connected layer in stage 1. Weights from the stage 1 encoder were transferred using transfer learning and the same tuned hyperparameters were applied for stage 2; stage 2 specific hyperparameters were tuned separately and used for final training. Because it is plausible that the risk captured by non-coding SNPs may affect a person’s risk observed in gene coding regions, we progressively unfroze the model’s layers during training, starting with only the stage 2 layers being unfrozen and eventually allowing all layers to be trainable.

In stage 3, a two headed model with adversarial learning via gradient reversal was created so that ancestry information could not predict disorder status (Ganin et al. 2016). For this, transfer learning was again applied to transfer learned weights from the stage 2 model and progressive unfreezing was enabled. Simultaneously, the 10 PC projections for each individual were used as labels and a reverse gradient layer was added to the adversarial head subsequent to the graph embeddings output. These embeddings were then entered into a linear layer and ultimately used to predict the ancestral labels with MSE loss serving as the loss function. In stage 3, the total loss is a combination of the main classification objective and the adversarial loss. In turn, the effect of the adversarial loss in total loss is modulated by the adversarial weight, λ, which requires careful tuning to prevent either insufficient debiasing or degradation of classifier performance. We implemented an adaptive scheduler that dynamically adjusted λ based on two competing objectives: (1) to minimize the R^2^ between predicted and true ancestry PCs (target R^2^ < 0.05) and (2) to maintain the main task AUROC above a minimum threshold. The scheduler was programmed to started with a 5 epoch long warm up phase, in which λ is linearly increased. If AUROC dropped more than 3%, λ was scaled by 0.9, and if more than 5%, it was scaled by 0.7; simultaneously, if R^2^ exceeded some specified critical value (here, 0.11), λ was increased by 1.25. If R^2^ was less than the critical value but above the target of 0.05, the adaptive scaler increased λ modestly by a factor of 1.08, and if the correlation between predicted and true ancestry was below the target, λ was slightly decayed to maintain stability. This arrangement precluded the model from using ancestral features and forced it to learn ancestry invariant patterns for predicting class status.

### GNN implementation and hyperparameter optimization

GNN models were developed using Pytorch version 2.7.0 + CUDA 12.8, torch-scatter torch-2.7.0+cu128, and PyTorch Geometric 2.7.0, in a containerized environment (Fey and Lenssen 2019; Paszke 2019). To address high computational demands, models were run using DistributedDataParallel with multiple GPUs; in this format, a copy of the model exists on each GPU, with each GPU receiving a minibatch of the dataset. After the backward pass, gradients are averaged across GPUs to ensure model stability. To further save memory during training, automatic mixed precision and gradient checkpointing were also used. Individual node features matrices, paired with phenotype and ancestry information were saved as Lightning Memory-Mapped Database (LMDB) files and input into the models alongside the adjacency matrices. To understand how a model optimized without any graph information might perform, we trained and tested a “Deep Sets” model, which ran the GAT model without any edge index; this resulted in the GAT behaving similarly to a multi-layer perceptron for each node.

Hyperparameter tuning was performed using Optuna with the objective of maximizing validation area under the curve (AUROC) (Akiba et al. 2019). 40 trials with 20 epochs each were conducted for stage 1, with early stopping at 10 epochs. For stage 2 and 3, 20 trials with 20 epochs were conducted, with the same early stopping rule. The following hyperparameters were optimized: dropout, number of GAT heads, hidden dimensions for the GAT layers, output dimensions for the GAT layer, learning rate, dimensions for the combined graph features and graph embeddings layer, lambda for the weight of the gradient reversal layer, and the weight of the adversarial task in computing the overall loss. Due to memory issues, different batch sizes were also dynamically evaluated for each Optuna trial, and the maximum size that did not yield an “out of memory” error was used. When performing the final training for the model, both the learning rate and batch size were scaled by 0.5 to prevent out of memory errors.

In addition to the GNN models, we evaluated polygenic risk scores (PRS) using generalized linear models (GLM) as a traditional genomics baseline via R 4.2’s base glm function. PRS were computed separately for whole-genome, intergenic, and genic variant sets. We used both individual Alzheimer’s disease PRS as well as combined scores incorporating PRS from genetically correlated disorders. For each PRS configuration, we fit a logistic regression model on the training set and evaluated predictive performance on the held-out test and validation sets.

### Model Performance Evaluation

Models were evaluated by computing test AUROC on the fully trained GNN models or PRS. To estimate the confidence interval of the area under the ROC curve (AUROC), we used the pROC package in R or a Numpy library implementation in python to apply non-parametric bootstrapping using number of bootstraps = 2000 and a confidence level of 95% (Harris et al. 2020; Robin et al. 2011). During each bootstrap iteration, stratified resampling was done to ensure class diversity. Cohen’s D, also implemented via Numpy, was used to quantify differences in degree between top and bottom nodes. Chi Squared testing was done using the scipy.stats package to assess prevalence of hubs amongst top nodes vs expected hub amounts. (Virtanen et al. 2020) The AUROCs between models were also compared using the pairwise DeLong’s test (DeLong et al. 1988).

To assess the correlation between the predicted log-odds (logits) of the GNN models and the PRS models, logits were standardized and plotted against each other; Spearman’s ρ and Pearson’s correlation coefficient were both calculated. Next, to investigate whether creating ensemble models improved classification performance, we constructed ensemble models using elastic net regularized logistic regression (EN) in R 4.2 via the glmnet and pROC packages (Friedman et al. 2010; Robin et al. 2011). For each base model (whole genome combined PRS and GNN), we extracted logits on the validation and test sets and normalized them. The EN ensembles were trained on validation logits using fivefold cross-validation to select the optimal regularization parameter. In these multiple predictor ensembles, we used EN with mixing parameter α = 0.5 (equal L1 and L2 penalty); the learned coefficients represent the relative contribution of each base model to the final prediction. As with individual models, performance was assessed on the held-out test set using AUROC, with 95% confidence intervals estimated via the DeLong method. To establish whether ensembles outperformed single classifiers, pairwise De Long’s test were again conducted.

### Post-hoc analyses and statistical testing

For all models using the BLC term, we extracted the BLC scale value and found the magnitude of the GAT and BLC term outputs after training was complete. Subsequently, we computed the cosine similarity between the GAT outputs and BLC output across the test sets and saved for later analysis.

To assess the importance of different nodes and edges with respect to model predictions, we performed a gradient-based analysis on the trained GNN, collecting different metrics depending on the objective. During this pass, we froze all model parameters and enabled gradient tracking only for the node features and graph-level covariates. For node level analysis, we computed gradients of the classification margin independently with respect to node level features (Simonyan et al. 2013). For the purpose of assessing cumulative importance, we took the mean of the signed gradients over each node’s input features and then obtained the absolute value of the mean. Subsequently, we summed these absolute values across all test samples to generate a final importance score for each node; nodes were then ranked by this score. We plotted cumulative importance by dividing the summed importance scores of the top-ranked percentage of nodes by the total importance score across all nodes. We obtained edge importance by taking the gradient of the model output with respect to the pre-softmax attention logit. To obtain cumulative importance for edges, the absolute value of edge importance, for every edge in the graph, was obtained for each sample and edges were ranked; edges were then ranked in an overall global list by average per sample rank and the sum of absolute edge importance was calculated on a per-edge basis. A Lorenz curve was plotted following the same approach as used for the nodes. Finally, to identify any differences that exist in the distribution of node or edge importance between GAT alone vs GAT + BLC term models, we computed several measures of importance distribution, including mean and minimum node/edge importance per stage, the bimodality coefficient, excess kurtosis, and the Gini coefficient.

To establish the most important nodes for ablation analysis, we took the mean of the signed gradients over each node’s input features and identified the 1635 nodes that had the highest importance values; we extracted this set of nodes for every sample in the validation set, and ranked nodes based on how frequently they appeared as highly important nodes for each sample – the 1635 nodes with the highest frequency were selected for ablation. Next, to identify the global least important nodes (bottom 10%), we took the absolute value of the node gradient per feature and obtained the mean value per node and extracted the 1652 least important nodes per validation set sample by this metric. The set of most consistently unimportant nodes was titled the global bottom 10%. The same process for finding the global top nodes was used to find the top edges; to obtain the global bottom edges, the absolute value of the attention weight gradients was obtained per edge and all edges ranked per-sample. The bottom 10% of edges for each sample were collected, and the least important edges by frequency in individual bottom-edge lists were deemed the global least important edges.

We performed three types of ablation analysis. In the first method, we removed either the top 10% of most important nodes/edges, the bottom 10% most important nodes/edges, and a random 10% subset of nodes and edges from the input dataset. Models were then subject to full retraining and the AUROC was assessed. We removed a random sampling of 10% of nodes or edges to distinguish the effect of removing the top scoring input features vs simply reducing graph or node feature information. Secondarily, we also used margin decomposition analysis to understand the relative contributions of either the GAT encoder or the graph level features to predictive performance. We hooked the fc fusion layer to capture actual per-sample activations, assessed encoder and graph feature activations, computed an effective margin vector, and obtained per-sample scalar cores for each component. Third, we assessed model predictions via inference by zeroing out either: all graph-level features, zeroing individual node or graph level features one at a time, or nullifying the GNN encoder output entirely. For this component-wise nullification analysis, we sought to assess whether feature contributions varied across the biomarker spectrum and subsequently stratified test samples into quartiles along three axes: (1) the mean node-level feature value across graph nodes, (2) the value of each individual graph-level feature, and (3) an encoder confidence score derived by projecting the GNN representation through the fusion layer weights and classification head, approximating each sample’s reliance on graph-topology information. Per-quartile AUROC was computed for the full model and each ablation condition, and the difference (AUROC drop) was reported as a measure of feature-pathway dependence within each stratum.

Finally, we sought to understand which components of the input data were operationalized to distinguish case vs control, and how the components used for class discrimination aligned with previous findings in the literature. For this, we performed GxI attribution analysis for individual node features, for each stage. We then computed Cohen’s D to quantify the effect of each feature on a per-gene basis and ranked the list of all genes in the graph based on effect size. To isolate genes with the greatest effects, z-scoring was performed on Cohen’s d values, and all genes ranked. This ranked list was subsequently used as input for a GSEA analysis via GSEApy (Fang et al. 2022). We also obtained FDR corrected p-values for each node via the Welch test and recorded any significant nodes, and looked for genes with concordance across features and multiplied their z-scores to quantify effects. We tested for enrichment among all pathways used in the graph that had a size greater than 15 but less than 199 genes; this restriction was put into place to maximize statistical power and prevent the dilution of signal. We then conducted additional enrichment analysis by comparing our ranked node list with pre-defined gene sets with specific functional properties. To do this, we used the fgsea package to analyze gene lists that corresponded to different properties; lists were obtained by querying enrichR, which accessed the following databases: (a) Brain atlases (Allen Brain Atlas up/down, Allen_Brain_Atlas_10x_scRNA_2021) to look for regional associations, (b) cell type markers (Azimuth_Cell_Types_2021, Tabula_Sapiens, PangaloDB_Augmented_2021, CellMarker_Augmented_2021, Descartes_Cell_Types_and_Tissue_2021) to identify cell type relationships (Cao et al. 2020; Franzén et al. 2019; Hao et al. 2021; Hawrylycz et al. 2012; Hu et al. 2023; Jones et al. 2022; Korotkevich et al. 2021; Kuleshov et al. 2016; Liberzon et al. 2015; Subramanian et al. 2005; Sunkin et al. 2013; Yao et al. 2021). fgsea permuted gene labels 10,000 times relative to the ranking statistic to build a null distribution of enrichment values that the running enrichment score is compared against to evaluate for enrichment; results were subject to false discovery rate correction within each functional gene set category and subsequently filtered for Brain relevance and use of human sourced gene sets(Korotkevich et al. 2021).

For edge analysis, given the large number of edges, we opted for a more focused approach and identified the subnetworks most relevant to AD case-control discrimination. For each diagnostic group we computed the mean importance score for every edge across all samples within that group, then retained the top 1% of edges ranked by group-level mean importance, yielding a group-specific subgraph. Selected edges were aggregated into an undirected graph, with bidirectional edges between the same gene pair counted once and connected components were extracted. Components were characterized by size, edge density, hub genes (top five nodes by degree centrality), and bridging genes (top five nodes by betweenness centrality). To assess the biological coherence of each subgraph, we performed hypergeometric pathway enrichment testing against the constituent pathways that the nodes mapped onto, with Benjamini–Hochberg correction applied at a false discovery rate threshold of 0.05. Finally, we compared case and control subgraphs by computing the Jaccard similarity of their node sets and performed separate enrichment analyses on the case-unique and control-unique nodes to identify pathways preferentially implicated in disease-associated versus baseline network activity.

## Results

Stage 1 GAT models yielded AUROCs ranging from 0.66, for the GAT with a bilinear context (BLC) term that ran using just AD gene scores (model no. 5), to 0.73, for GAT with a bilinear module operating with AD and correlated disorder gene scores (model no. 2) (Table 1a). Stage 1 Deep Sets models using node features from the Pathway Graph, both with and without BLC had AUROCs of 0.70 (model nos. 3 and 4). This result did not diverge significantly from the predictions of most other non-Deep Sets models, with the exception of models 2, 5, and 7. Varying edge density in the Hippocampus gene expression graph for GAT models without the BLC term did not substantially affect AUROC (model nos. 6, 9, and 10), and neither did the use of an AD patient gene expression graph vs control (model nos. 6 and 8). Use of the bilinear context module did not change affect AUROC in deep-sets mode with Pathway Graph features (model nos. 3 and 4), but it did significantly worsen performance compared to the GAT when using the 0.25% Hippocampus control graph (model nos. 6 and 7). Use of the BLC term alongside the Pathway Graph significantly improved AUROC compared to the Pathway Graph GAT alone and Deep Sets mode (model nos. 1, 2, 3) as well as all gene expression derived models.

**Table 1a:**
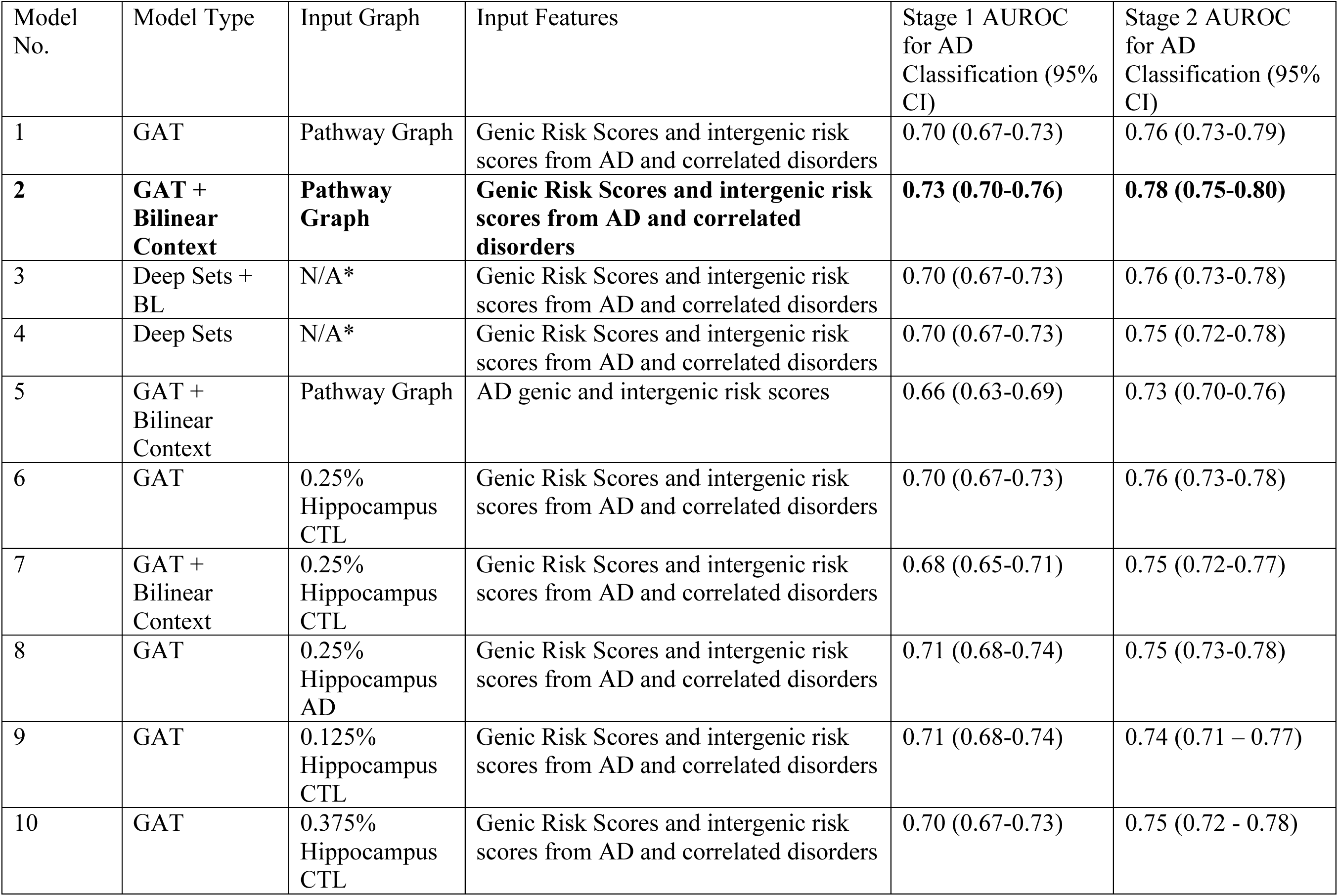
Stage 1 and Stage 2 AUROC for AD Classification Across GNN Model Configuration.

Stage 2 models showed similar trends to Stage 1. All Gene Expression Models, regardless of edge density or source phenotype yielded AUROCs of around 0.74-0.76 and were not significantly different from one another (model nos. 6, 8, 9, 10). Both Pathway Graph Deep Sets models, with or without BLC, (model nos. 3 and 4) showed similar AUROCs, also of 0.75-0.76, and were not significantly different from each other, or from the GAT Pathway Graph model (model no. 1). The GAT with bilinear context module operating on the Hippocampus 0.25% control graph (model no.6) recovered from a poor stage 1, yielding AUROC of 0.75, but the single feature Pathway Graph (model no. 5) with bilinear context was significantly worse than the other models, with an AUROC of 0.73. As before, the GAT Pathway Graph with correlated disorders and bilinear context (model no. 2) was significantly better at classification than all other models, with an AUROC of 0.78. All models improved significantly from stage 1, with an increase in AUROC ranging from 0.03 to 0.07, and an average boost of 0.048. Altogether, node and edge importance distributions showed distinct patterns across training stages, and across model architecture; results for GAT (model no. 1) and GAT-BLC (model no. 2) are presented in Supplementary Table 3.

Stage 1 dual head models showed that models did not significantly predict ancestry, even without any debiasing. PC correlations for all tested models (models no. 1, 2, 3, 4, 5, 6) were below zero, and adversarial MSE was above 1.0. Debiased stage 3 models were largely aligned with their stage 2 predecessors, with most model AUROCs being within 0.01 of the original model; stage 1 models 3 and 5, e.g. Deep Sets with BLC and Hippocampus 0.25% GAT showed a slight improvement of 0.01 (Supplementary Table 2). As in the dual-head, R^2^ was less than 0; MSE was at least 0.95 in all models.

The most important hyperparameters for the Deep Sets GAT BLC model (model no. 3) were: dropout, FCNN2 size, and weight decay, with a relative importance of 0.43, 0.23, and 0.12, respectively. In stage 2, they were learning rate (0.51), batch size (0.17), and dropout (0.14). In contrast, for the GAT BLC Pathway Graph (model no. 2), the top parameters were FCNN2 size (0.23), learning rate (0.21), and output dimensions (0.17). In stage 2, it was dropout (0.34), freezing of the FC GNN layer (0.34) and batch size (0.21). Bilinear scale for model 2 ranged from −0.01 in stage 1 to 0.01 in stage 2, while remaining stable at 0.11 for model 3 (Supplementary Table 1). Model 2’s GAT and BLC outputs were nearly orthogonal and of the same magnitude, but in model 3, the Deep Sets output and BLC showed a strong negative correlation of −0.59 and an exploding magnitude for BLC. In model 7, where the BLC term actively hurt performance, the bilinear scale output was roughly 0.08, cosine similarity 0.30, and the magnitude around 10X higher than the GAT output.

All GNN models we trained were better than chance (AUROC = 0.5) at classifying AD (Table 1a). They were not, however, by themselves, more accurate than the PRS linear models, which yielded identical results using just AD and AD and the correlated phenotypes. Our best GNN model had an AUROC of 0.78 (0.75-0.80), compared with 0.80 (0.78-0.83) for Whole Genome PRS (Table 1b). Our ensemble models of stage 1 and the PRS were non-significant; however, the stage 2 and 3 ensembles (AUROC of 0.82) yielded significantly better performance than PRS alone (0.80). Logits of model 2 for each stage were plotted against logits for the whole genome PRS model, yielding r = 0.58 for stage 1, with an increase to r = 0.69 in stage 2, and a subsequent drop to 0.66 in stage 3 (Figure 4a). Ensemble elastic net models of PRS with the different stages showed slightly greater coefficient magnitudes for the whole genome PRS relative to the GNN models (Figure 4b). Altogether, stage 2 and 3 + PRS ensembles (AUROC of 0.82) were statistically better than whole genome PRS alone (0.80) (Table 1b).

**Table 1b:** AUROC for PRS and PRS ensemble models.

| Model No. | Model Type | Input Features | AUROC for AD Classification (95% CI) |
| --- | --- | --- | --- |
| 12 | Logistic Regression | Whole Genome PRS from AD and correlated phenotypes | 0.80 (0.78-0.83) |
| 13 | Logistic Regression | Whole Genome PRS from AD | 0.80 (0.78-0.83) |
| 14 | Logistic Regression | Intergenic PRS from AD and correlated phenotypes | 0.70 (0.67-0.73) |
| 15 | Logistic Regression | Intergenic PRS from AD | 0.70 (0.67-0.73) |
| 16 | Logistic Regression | Genic PRS from AD and correlated phenotypes | 0.78 (0.75-0.80) |
| 17 | Logistic Regression | Genic PRS from AD | 0.78 (0.75-0.80) |
| 18 | Elastic Net | Logits from Model 12 and Stage 1 of Model 2 | 0.81 (0.78-0.84) |
| **19** | **Elastic Net** | **Logits from Model 12 and Stage 2 of Model 2** | **0.82 (0.79-0.84)** |
| **20** | **Elastic Net** | **Logits from Model 12 and Stage 3 of Model 2** | **0.82 (0.79-0.84)** |

Cross-sample consistency analysis revealed that a small proportion of nodes were reliably important across test samples, while edges showed markedly less consistency (Figure 2). At the >50% threshold (i.e. nodes ranked as more important than half of the test set), 14.2% and 11.4% of nodes in Stage 1 and 2 exceeded this criterion. This proportion declined steeply at more stringent thresholds; only 1.1% and 0.5% of nodes remained at 90% consistency. Stage 1 exhibited higher node consistency than Stage 2 across all thresholds. Edge cross-sample consistency was substantially lower, with only 7.25 % and 6.73 % of edges exceeding the >50% threshold in Stage 1 and 2, respectively, and 0.45% and 0.25% at >90%. Lorenz curves showed that the top edges and nodes in both stages explain much more importance than would be expected if all input graph components were equally important. Altogether, importance was more concentrated in top edges than the top nodes for both stages.

**Figure 2:**
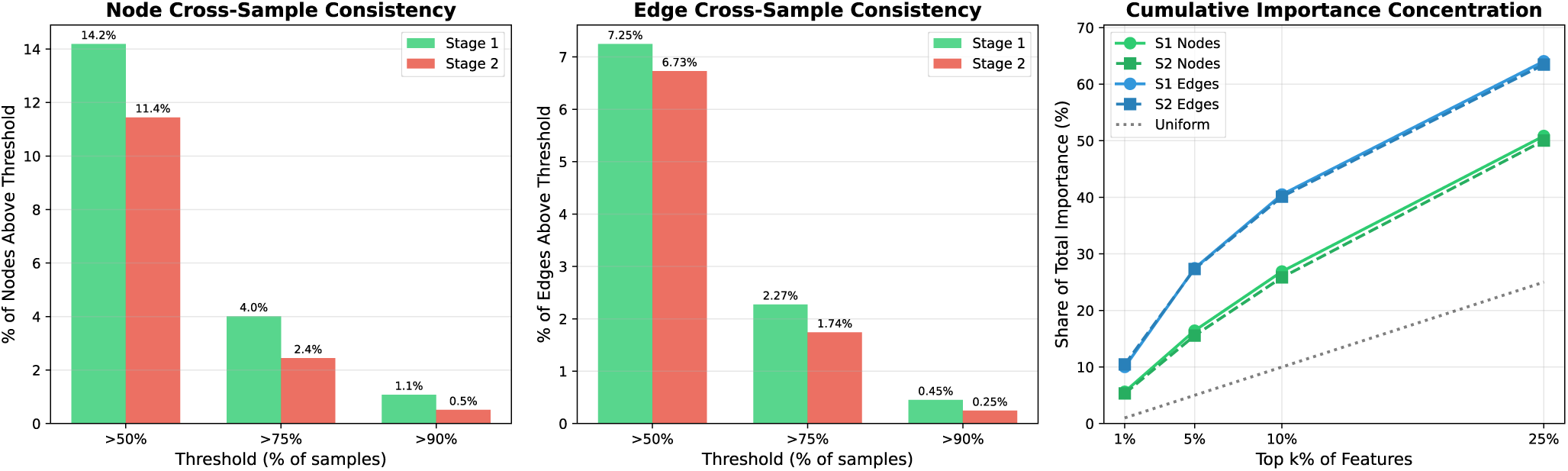
Explainability Analysis: Importance Distribution and Cross-Sample Stability

**Figure 3:**
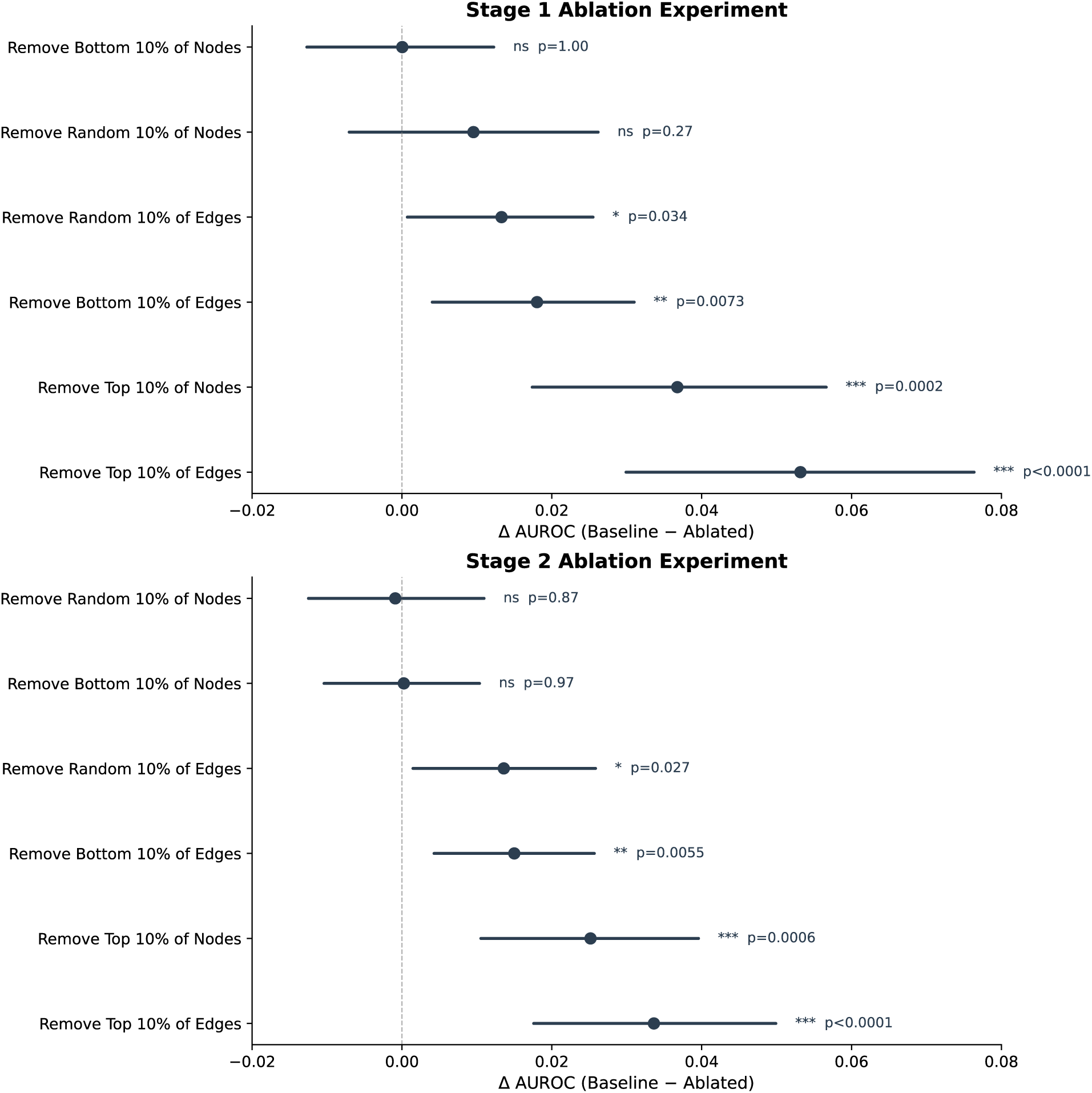
Ablation Study: Impact of Node and Edge Removal on AD Classification Performance

Node ablation models had varying effects on the number of edges and average node connectivity (Supplementary Table 4a-c). Removing the bottom nodes resulted in a loss of just 5% of edges and raised the average overall node degree for the resulting graph, while removing a random set of 1635 nodes led to the disappearance of 18.3% of the graph’s edges and lowered the average degree from 57.1 to 52.2 (Supplementary Table 4a). Depletion of the top nodes yielded the largest effect, with the loss of 28% of edges and a new average degree of 45.9 (Supplementary Table 4b). Subsequently, the top 10% most important nodes were found to have a 15X higher degree than the bottom 10%, with Cohen’s D of 1.72 (p<0.001) and around a 1.55X higher degree than a random set of nodes (Cohen’s D of 0.46, p <0.001). Top edges were shown to be connected to be top nodes more than expected by chance, with approximately 31.4% of top edges in stage 1 and 32.9% of top edges in stage 2, connecting a top node, and roughly 3.13% and 4.09% are connections between top nodes (chi-squared, p <0.001). The effects are magnified at more stringent thresholds; at 1%, the odds ratio (OR) of a top edge connecting to a top node is 1.66 and 1.55 for stage 1 and 2, respectively, (relative to 1.20 and 1.26 at 10%), and connections between top nodes have ORs of 2.12 and 1.95 (versus 1.11 and 1.24); all ORs are significant. Conversely, the top 10% of nodes by importance were found to be significantly under-enriched for hubs via Chi Squared (p<0.001) and do not have more top edges than expected by chance. Additionally, removal of the top 10% of edges resulted in approximately 100x more isolated nodes and more than 3x more single degree nodes compared to removal of a random subset of edges; removal of bottom edges had little effect on either metric (Table 3c). Removal of the top 10% of nodes resulted in a significantly larger in AUROC vs removal of random or the bottom 10% of nodes in both stages (Figure 2a-b). Surprisingly, removal of the any subset of edges in stage 1 significantly reduced performance relative to baseline, with the worst result being seen with removal of the top 10% of edges, which reduced AUROC to 0.68 (Figure 2a, Supplementary Table 4c). In stage 2, this trend persisted, albeit the drop in performance for the top 10% edge ablation graph was somewhat reduced relative to stage 1 (Figure 2b, Supplementary Table 4c).

Under component-wise nullification, the encoder-only condition (graph features nullified) achieved an AUROC of 0.74, the graph-feature-only condition (encoder nullified) achieved 0.70, and the intact model achieved 0.78. Encoder activations exhibited substantially larger magnitude (mean = −3.10, SD = 1.46) compared to graph-feature activations (mean = −0.002, SD = 1.11), and the encoder produced a greater case–control separation (difference = +1.21) than graph features alone (difference = +0.76). In tandem, zeroing the GNN encoder output while retaining all graph-level features reduced AUROC from 0.78 to 0.70 (drop = 0.08), whereas zeroing graph-level features while retaining the encoder reduced AUROC to 0.74 (drop = 0.04). Zeroing individual graph level features had little effect on AUROC, except for AD intergenic PRS, which yielded a nearly 0.04 AUROC decline. Simultaneously, elimination of the AD node level feature was also damaging in stage 2, reducing AUROC to 0.63, a greater effect than when Schizophrenia genic scores were removed (AUROC = 0.69) (Figure 4). However, zeroing the fluid intelligence node feature channel was by far the most damaging in stage 2, reducing AUROC to 0.50. In Stage 1, AD was the most critical node feature channel (AUROC = 0.50 upon ablation), followed by fluid intelligence (AUROC = 0.63 upon ablation) and putamen volume (AUROC = 0.65 upon ablation).

**Figure 4:**
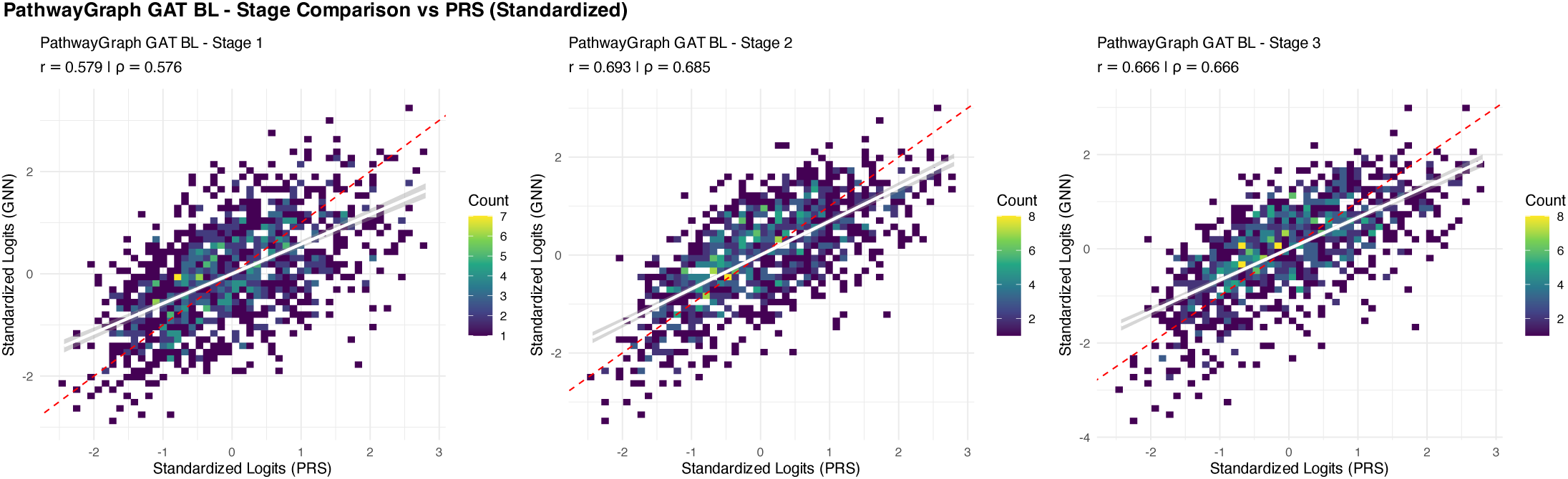
Correlation Between GNN and PRS Logits Across Training Stages“

When the encoder was highly confident (Q4, mean score = −1.3), zeroing all graph features reduced AUROC by only 0.030, whereas for the three lower quartiles the reduction ranged from 0.12 to 0.13. Individual feature ablation within these strata confirmed that AD PRS was the only graph-level feature with individually meaningful impact: its ablation reduced AUROC by +0.11 to +0.15 in Q1–Q3 but only +0.03 in Q4; all remaining 11 graph features had negligible individual ablation effects (<0.01 in every quartile), indicating that their contributions are either mutually redundant or too small to detect in isolation (Supplementary Figure 1b). For AD node level features in Stage 2, the graph-level AD intergenic feature was most impactful when the node-level AD signal was in the middle range (Q2 drop = +0.09) and least impactful at the high extreme (Q4 drop = +0.02). In contrast, for Fluid Intelligence node level features, the graph-level AD intergenic feature had the largest effect in Q4 (+0.07), with reduced effects in Q1 (+0.05) and Q1 (+0.05), and no effect on Q3 (Supplementary Figure 1c).

In Stage 1, the AD genic attribution distribution was centered near zero (mean = 0.003, median = 0.003, SD = 0.062), with 52% of genes showing positive and 48% negative centered effect sizes, indicating that the base encoder distributes AD PRS signal broadly and without strong directional bias across the genome (Supplementary Figure 2). In Stage 2, following fusion training, this distribution shifted dramatically: the mean centered Cohen’s d rose to 0.30 (median = 0.32, SD = 0.09), with 100% of genes showing positive values. Fluid intelligence showed a complementary pattern: in Stage 1, its distribution was nearly symmetric around zero (mean = −0.01, SD = 0.06), but in Stage 2, the distribution shifted strongly negative (mean = −0.27, median = −0.29, SD = 0.10), with 98% of genes showing negative centered effect sizes.

Four genes survived FDR correction for individual significance in stage 1 across the AD and FI node feature channels: APOE, TOMM40, APOA2, and AFM; 3 others reached individual significance across at least one of the remaining 10 node feature channels (APOC1, APOA1, and LRIT3). In stage 2, 15, 452 and 14, 731 genes reached individual level significance for AD and FI, respectively, with an overlap of 14, 145 genes; however, all but 22 of the overlapping genes had opposing directionality (Supplementary File 1, Supplementary Table 5). Using z-score ranking, GSEA analysis among pathways used in constructing the graph yielded two significant pathways for stage 1 in FI: Plasma lipoprotein remodelling (NES = −2.28), and Neddylation (NES = −2.17) and seven pathways for stage 2: Potassium Channels (NES = −2.57, FDR <0.001), Collagen chain trimerization (NES = −2.31, FDR <0.01), Voltage-gated K+ channels (NES=−2.14), Cell-Cell communication (NES=−2.18), Amino/nucleotide sugar metabolism (NES=−2.14), Cell junction organization (NES=−2.15), and Neuromuscular process controlling balance (NES = −2.19); FDR was <0.05 for the latter 5 pathways. No pathways were significant for the AD node feature channel in stage 1, while six were in stage 2: MET activates PTK2 signaling (NES= +2.33, FDR <0.01), Collagen chain trimerization (NES = +2.3, FDR <0.01), Cardiac conduction (NES = +2.16), Collagen degradation (NES=+2.18), G alpha signalling (+2.09), SLC neurotransmitter transport (NES = +2.09); FDR was <0.05 for the latter 4 pathways.

Functional GSEA on z-score ordered lists resulted in 19 significant hits for the FI channel and 84 for AD for stage 2 (Supplementary File 1). The strongest signal for AD comes from enrichment in brain cell type gene sets, particularly deep-layer inhibitory neurons e.g. the LAMP5 CRABP1 interneuron set in layers 5–6 has the lowest p-value (p = 1.63×10⁻⁸), and several PVALB subtypes and VIP interneurons (IGDCC3, BSPRY) also show strong enrichment. At the same time, excitatory neuron sets from layers 3–6, including FEZF2 subtypes, are similarly enriched in the disease-enriched direction. Cell marker gene sets also reinforced a broad neuronal and glial theme — the Lake Ex3 excitatory marker set, oligodendrocyte precursor markers, Schwann/satellite glial sets, and Bergmann glia all reached genome-wide significances. Additionally, an immune signal also emerged, with two microglia et and an M1 macrophage set enriched in opposing directions. In contrast, every significant pathway in the fluid intelligence feature was enriched in the control direction, with the strongest signals originating from pyramidal cell markers and the Lake Ex3 excitatory neuron set, followed by oligodendrocyte precursors, GABAergic and glutamatergic neuron sets, and enteric neurons. Astrocyte/Schwann/Bergmann glia and satellite glial cell sets are all depleted for FI, and notably the microglia set (TYROBP CD74) that was positively enriched for AD is here negatively enriched.

Two subgraphs were generated in Stage 1 for case and control, comprised of 9, 387 edges mapping onto 7,505 and 7,509 genes, respectively (Supplementary File 2). While both subgraphs were dominated by a single large connected component: 5,882 nodes in case and 5,930 in control, the remaining genes were also distributed in smaller components that corresponded to well defined biological modules, including a nuclear pore complex module, histone/chromatin complexes, MAPK/PI3K signaling, and others. The subgraphs for both diagnostic groups were very similar (Jaccard = 0.941) and shared 7,280 genes; only 225 and 229 were unique to the case and control subgraphs, respectively. Accordingly, high concordance was also observed in hub genes (defined via degree centrality). However, enrichment analysis for the class unique genes identified some distinct signatures; case-unique genes were enriched for membrane protein ectodomain proteolysis (NES = 15.6, p<0.05), and amyloid fiber formation (NES = 8.7, p<0.05), while control genes were enriched for neuron maturation (NES = 17.9, p < 0.01), multivesicular body sorting (p<0.05), and glycolysis/energy reserve metabolism (p<0.05).

The stage 2 case and control subgraphs contained 7,666 and 7,664 genes, respectively, and were again dominated by a single large component 6,219 nodes in cases, 6,241 in controls). However, the subgraphs showed substantially greater divergence than in Stage 1; Jaccard similarity declined to 0.869, and there were 539 case-unique and 537 control-unique genes, more than twice the amount in the prior stage. Hub genes also differed; case-preferring hubs genes included PTPRU, APOBEC3A, NEBL, and HOXB5, while control-preferring hubs included TMEM33, USH1G, SDF2, GTF2F2, and SHCBP1L. Smaller components partially overlapped with those from stage 1 but demonstrated some reorganization; the nuclear pore subgraph grew to 33 nodes in the case subgraph, while the control held only 20 genes. Hypergeometric pathway enrichment analysis of the subgraphs revealed unique signatures for each diagnostic group. The control subgraph was enriched for response to unfolded protein (p < 0.001, NES = 1.89), ER unfolded protein response (p < 0.001, NES = 2.00), and Toll-like receptor cascades (p < 0.001, NES = 1.61), while the case subgraph showed unique enrichment for cellular response to heat stress (p < 0.001, NES = 1.95) and cellular response to metal ion (p < 0.001, NES = 1.85). Analysis of specific case and control-unique genes reinforced subgraph enrichment results; the most significantly case-enriched pathway was protein maturation by iron-sulfur cluster transfer (p<0.001, NES=16.5), with sub-pathways for [4Fe-4S] (p<0.001, NES=17.3), and [2Fe-2S] cluster transfer (p<0.001, NES=22.8), also being significantly enriched. Additional case-unique enrichments included pyrimidine nucleotide biosynthesis (p<0.01, NES=7.14), negative regulation of Wnt signaling (p<0.01, NES=3.93), nuclear pore assembly, and human cytomegalovirus late events (p<0.01, NES=4.74). In contrast, the stage 2 unique control genes were enriched for ER calcium homeostasis (p<0.001, NES=30.5), synaptonemal complex assembly (p<0.001, NES=10.2), protein insertion into ER membrane (p<0.001, NES=7.61), and mitochondrial transport (p<0.001, NES=3.26),

## Discussion

Logits correlation analysis show that while our best performing GNN and PRS models yield somewhat similar AUROCs of 0.78 and 0.80, respectively, the amount of shared variance is not high. As a result, ensemble models comprised of stage 2 or 3 and the PRS performed significantly better (AUROC of 0.82) than any of the PRS models. This indicates that the GNN is extracting useful information, although the absolute increase in AUROC (∼2.5%) is not dramatic. Our results thus show that the additional context of gene-gene associations provided by a graph improves the classification of AD from GWAS data. However, this context does not appear to be used effectively by GAT models without the addition of the BLC term as there was no significant difference between models that had no context at all (Deep Sets) vs different GNN configurations without the BLC term, suggesting the node features alone carry significant signal. The best performing model uses the Pathway Graph defined from a priori known pathways and some Ricci rewiring in conjunction with the BLC term.

The intent of the BLC term was to approximate global context by creating a global representation vector and finding the effect of global context on each node’s (local context informed) representation of genetic risk by multiplying the two vectors. In turn, this product is then reprojected to match the dimensions of the original node representation and added in a scaled fashion to the aforementioned representation. Given that the bilinear scale initializes at 0.1 and ends at −0.01 for stage 1 training and only inches forward in stage 2 to 0.01 for the best performing model, and the fact that the GAT and BLC outputs are effectively orthogonal, it is unlikely that there is a direct representational contribution from interactions between global genetic risk and individual node risk in the trained model. In contrast, worse performing models that also incorporated the BLC term, e.g. Deep Sets and the Hippocampus Gene Expression GNN models both had non-zero bilinear scale values, greater magnitude differentials between the GAT and BLC output (10 to 100x), and non-zero cosine similarity between the two outputs. Intriguingly, the Deep Sets model with the BLC term, which has no performance advantage over the Deep Sets model without the BLC term, has negatively correlated GAT and BLC outputs, while the Hippocampus GE + BLC model, which actively underperformed its non-BLC counterpart, showed a positive correlation.

Because GNN models without the BLC term, regardless of graph type, yield the same classification performance, we propose that the BLC works by amplifying the use of graph structure during training. Getting orthogonal outputs requires either the focused depression of the BLC term by creating less useful Wu and Wv matrices and lowering the bilinear scale, or by diminishing the BLC term in conjunction with backpropagation to the GAT layers. The former mechanism should not change AUROC, but we do see shifts for both the Hippocampus GE + BLC and Pathway Graph with BLC. Subsequently, we argue that for the Pathway graph model, the BLC computed global context is incorporated into early graph embeddings; however, as training progresses, the model finds the global context to be unhelpful. In response, the model actively supresses the nodes and edges that provide said context, resulting in improved GAT performance. In contrast, gene co-expression networks are known to be noisy, and as a result, the Hippocampus GE graph likely has fewer reliable edges. Consequently, the updated node representations after the GAT layers are noisy, and so too, is the global context computed by the BLC term. However, since the input graph structure itself is noisy, backpropagation cannot facilitate the de-prioritization of problematic edges, leading to worsened performance. As a result, the final bilinear term in the trained model is non-zero and correlated with the GAT outputs, yielding worsened performance. The Deep Sets model has no graph structure, so the BLC term may fill a structural void, leading to an exploding magnitude (866X). Simultaneously, the model also has greater flexibility and can compensate through the main prediction, yielding the negative cosine similarity value.

The difference in distribution of node and edge importance across the baseline Pathway Graph GAT (model no. 1) vs Pathway Graph GAT BL (model no. 2) is consistent with this argument. Node Gini coefficients are similar across models in both stages, indicating comparable concentration of feature importance, but there is greater variance in model 2, as demonstrated by higher kurtosis effects for both edges and nodes, and a bimodality coefficient of 0.64, which indicates a clear separation between a small set of highly important nodes and another set with near zero importance. This suggests that the GAT + BLC model learns to differentially weight specific input nodes and edges on the graph while the simple GAT does not, perhaps because the node features are already rich in predictive information. This is reinforced when the changes in stage 1 and 2 importance metrics are compared between models; in model no. 1, node kurtosis shifts only modestly from 1,080 to 1,062, bimodality coefficients are stable, and Gini coefficients decrease marginally, with edge importance following a similar pattern of relative stability. Conversely, in model no. 2, a bigger shift is visible; node kurtosis drops from 6,393 to 330, bimodality falls from 0.64 to 0.32, and maximum node importance decreases from 0.20 to 0.031, while edge kurtosis quadruples. The model appears to shift its representational strategy: from a node-concentrated, edge-distributed pattern in Stage 1 to a node-distributed, edge-concentrated pattern in Stage 2. When intergenic PRS features are introduced through the fusion architecture, the model no longer depends on a small number of critical pathway hub genes, as it now has access to complementary genomic information that allows node importance to be distributed more broadly. This cross-stage reorganization does not occur in the baseline GAT, which lacks the mechanism to modulate how node features interact with graph topology. The results of ablation experiments align with this, showing significantly reduced AUC when the top nodes are removed but no difference with the bottom nodes or removal of random nodes. The strong effect of node removal is likely due to the changes in graph connectivity as well as the loss of important input features.

Removal of correlated phenotypes as node features (model no. 5) also significantly reduced performance. This phenomenon may be fueled by the fact that the node hidden representations formed after the GAT layers are less rich and relatedly, because the learnable attention parameter is computed based off the node features (equation 1a). Altogether, component-wise nullification experiments showed that the model relies more heavily upon graph representations than graph features, with channel-level ablation identifying AD and fluid intelligence genic risk as the dominant node-level features driving performance. Our findings show that the model relies on graph-level AD intergenic scores mostly when the encoder representation is less confident, consistent with a fusion strategy in which the intergenic PRS compensates for uncertainty in the pathway-level signal rather than contributing uniformly. At the same time, cross-stratification of graph-level AD feature importance by node-level feature quartiles also seems to indicate that the model uses intergenic AD signal differently depending on the node-level context.

The largest reduction in performance among the ablation models that underwent full training and optimization was observed on graphs in which either the top 10% most important edges or the top 10% most important nodes were removed. This is consistent with what one would expect for features that the model relies upon most heavily. However, significant performance reductions were also seen when depleting either a random set of edges, or the bottom 10% of edges. At first glance, the results are surprising, but disruptions in connectivity may impair message passing regardless of edge importance. This argument reinforces the idea of noise as a cause for underperformance in gene expression graphs – unreliable edges can greatly disrupt graph structure. Simultaneously, the decline in performance observed upon removal of random edges may also be the result of limited consensus amongst top edges; only 6-7% of edges are present in the top edges per sample for at least 50% of the samples.

Despite stage 2 (and 3) logits being more strongly correlated with whole genome PRS model logits than stage 1, ensemble models comprised of stage 2 or 3 and the PRS were significant while the stage 1 and PRS model was not. This suggests that stage 1 may reconstruct additive risk while stage 2 retains additive risk and adds additional orthogonal signal. Though stage 1’s architecture explicitly incorporates local epistasis, it is possible that the prediction is largely dominated by additive risk; this is consistent with challenges in detecting epistasis in other approaches (Balvert et al. 2024; Chatelain et al. 2018). Nonetheless, there are a few plausible mechanisms for the discrepancy between stage 1 and 2 ensembles: first, the integration of intergenic features may provide useful regulatory context that amplifies genes with limited coding-region signal but substantial non-coding involvement – this is consistent with previous studies showing that significant GWAS heritability lies in noncoding regions. Indeed, we know that in stage 2, large shifts in Cohen’s D directionality are observed for different node channels; thus, the context provided by intergenic features may push the model to learn differential usage of the node features, allowing for enhanced class discrimination. In particular, given that addition of intergenic features yielded opposing directional shifts for most genes (>14,000) across both the AD and Fluid Intelligence phenotypes, with only 22 genes having concordant directions, it appears that the model has learned the two phenotypes represent risk in opposite directions, which is biologically consistent. Second, as graph-level features partially capture aggregate risk, this may “free” the model to focus on more subtle gene level signals that were previously overshadowed by main effects. Importance metrics from stage 1 and 2 indicate dramatic reductions in average importance for both nodes and edges, with the effects being more pronounced for edges. This could result in more patient specific patterns of edge importance which ultimately reduce consensus importance. Given that attention weights are dependent on node features, it is reasonable for there to be heterogeneity in edge importance.

Evaluation of significant genes and pathways obtained via our analyses shows high concordance with known AD signal. The four genes that survived FDR after case-control importance comparison: APOE, TOMM40, APOA2, AFM have all been strongly linked to AD; in particular, TOMM40 is in tight linkage disequilibrium with APOE and has been implicated in age-of-onset modulation in AD(Chiba-Falek et al. 2018; Roses et al. 2010; Roses et al. 2013). Functional GSEA revealed that AD attribution concentrates in deep-layer inhibitory interneuron and VIP interneuron marker sets, independently recovering the same cell-type vulnerability hierarchy identified by single-nucleus transcriptomic studies of postmortem AD brains, but here via genetic risk(Consens et al. 2022; Mathys et al. 2024). The 22 concordant genes seem to segregate into three groups: “case-driven” and “AD negative, FI positive”, and “control.” In the former group, APOE is again recapitulated, but so were other AD hits such as EGR1, an immediate early gene critical for synaptic plasticity and memory consolidation(Koldamova et al. 2014). Finally, extracted subgraphs also capture multiple well characterized AD pathways, including stage 1 case-unique enrichment of amyloid fiber formation and membrane protein ectodomain proteolysis and stage 2 unique enrichment for iron-sulfur cluster transfer and metallothionein genes, which may align with the metal dyshomeostasis hypothesis of AD(Braymer and Lill 2017; Jassal et al. 2020; Myhre et al. 2013; Saftig and Lichtenthaler 2015). In contrast, control subgraphs in both stages seem to represent homeostatic and quality control mechanisms, including neuron maturation (stage 1) as well as ER calcium homeostasis and the unfolded protein response (stage 2).

Beyond validating established AD biology, some findings may represent novel insights from the model’s attribution architecture. The emergence of MET/PTK2 signaling as the strongest AD PRS pathway implicates hepatocyte growth factor receptor signaling in AD genetic risk; though this pathway has been explored in neuroinflammation and microglial phagocytosis, it is underexplored in the context of AD(Desole et al. 2021; Wright and Harding 2015). In tandem, potassium channels as the dominant fluid intelligence pathway may suggest that ion channel infrastructure supporting neuronal excitability may play a greater role in the genetic basis of cognitive resilience than previously appreciated(Alam et al. 2023; Kessi et al. 2020). Finally, the case-driven concordant gene KDM2A, a histone demethylase involved in chromatin remodeling, emerges as the second-strongest z-product after APOE, potentially reflecting a shared epigenetic substrate linking AD risk and lower fluid intelligence(Santana et al. 2023; Tsukada et al. 2006).

There are several limitations to our study. First, our sample size is relatively limited, with only 7358 participants. This reduces statistical power and can lead to greater variance; reproducing this result on a larger external sample or using transfer learning from a model trained on a larger sample could help. Another possible challenge is class imbalance; although we sought to address this by using class weights, it may have helped to use other approaches, such as oversampling the minority class. Samples also came from seven different studies, which could have caused a batch effect; use of study as a covariate when creating the train, test, and validation splits would address this. Another challenge is the use of additive genetic scores as input. This inherently limits our ability to look for interactions between loci given that the risk within a physical region has already been reduced to a sum. An alternative approach could be using raw genotypic data, with genic SNPs serving as node features. Due to time constraints, we were also only able to run analyses on a single split of data; running five-fold cross validation from a combined validation and training set while maintaining a held-out test et could reduce variability and increase robustness.

There are multiple avenues for development in terms of model architecture. In this work, we only used undirected edges; incorporated directed edges could add useful regulatory information about relationships. We also did not take advantage of edge features, which could have helped distinguish between different types of networks or relationships between the genes, e.g. transcription factor to target gene, inflammatory pathways vs metabolic pathways, an edge for physical proximity on the chromosome versus functional connection etc. Another potentially useful approach could be the use of different types of nodes to signify gene characteristics, e.g. some genes may be preferentially expressed in certain cellular compartments, developmental stages etc. We also injected intergenic PRS as graph level features; it could be useful to decompose the scores into functional categories like enhancers or promoters. Non-coding risk loci could also be incorporated into the graph structure via heterogeneity in node type. Lastly, it may be useful to use a dataset with multiple phenotypes, e.g. mild cognitive impairment, mixed dementia, or traumatic brain injury to assess the specificity of the predictions.

Altogether, our results show that deploying interpretable GNN models on graph structured GWAS data provides useful information for prediction that is orthogonal to additive PRS models and modestly improves classification performance. We further find that these GNN models are highly sensitive to graph structure and perform better on graphs comprised of well characterized interactions. To the best of our knowledge, this is the first use of GAT layers to learn representations of genome wide genetic risk informed by local epistasis for the purposes of classifying AD; previous work with GNNs in the context of AD limited genetic risk to missense variants, only used PPIs vs established pathways, and did not consider PRS or correlated phenotypes (Hernández-Lorenzo et al. 2022). In tandem, this approach facilitates the generation of hypotheses about genetic contributions to risk of AD development, including interactions between genomic loci.

---

**Supplementary Table 1:** Stage 3 AUROC and PC prediction metrics for different GNN configurations.

| Model No. | Model Type | Input Graph | Input Features | Stage 3 AUROC for AD Classification (95% CI) | Estimated PC MSE | Estimated PC R2 |
| --- | --- | --- | --- | --- | --- | --- |
| 1 | GAT | Pathway Graph | Genic Risk Scores and intergenic risk scores from AD and correlated disorders | 0.75 (0.72-0.77) | 1.6 | < 0 |
| 2 | GAT + Bilinear Context | Pathway Graph | Genic Risk Scores and intergenic risk scores from AD and correlated disorders | 0.77 (0.74-0.80) | 0.95 | < 0 |
| 3 | Deep Sets + BL | N/A* | Genic Risk Scores and intergenic risk scores from AD and correlated disorders | 0.76 (0.73-0.79) | 1.6 | < 0 |
| 4 | Deep Sets | N/A* | Genic Risk Scores and intergenic risk scores from AD and correlated disorders | 0.76 (0.73-0.78) | 1.8 | < 0 |
| 5 | GAT + Bilinear Context | Pathway Graph | AD genic and intergenic risk scores | 0.73 (0.70-0.76) | 2.2 | < 0 |
| 6 | GAT | 0.25% Hippocampus CTL | Genic Risk Scores and intergenic risk scores from AD and correlated disorders | 0.76 (0.74-0.79) | 3.5×10<sup>3</sup> | < 0 |

**Supplementary Table 2:** Metrics for Bilinear Models.

| Model | Stage | Bilinear Scale | Cosine Similarity | Cosine Similarity Std | GAT Output Magnitude | BLC Output Magnitude |
| --- | --- | --- | --- | --- | --- | --- |
| Pathway Graph BL | 1 | -0.01 | 0 | 0.02 | 2.7 | 3.2 |
| Pathway Graph BL | 2 | 0.01 | 0.02 | 0.01 | 2.6 | 2.6 |
| Deep Sets | 1 | 0.11 | -0.59 | 0.02 | 7.5 | $8.7 \times 10^2$ |
| Deep Sets | 2 | 0.11 | -0.59 | 0.02 | 7.5 | $8.7 \times 10^2$ |
| Hippocampus<br>CTL 0.25% | 1 | 0.08 | 0.30 | $3.0 \times 10^{-3}$ | 0.60 | 6.8 |
| Hippocampus<br>CTL 0.25% | 2 | 0.08 | 0.30 | $3.0 \times 10^{-3}$ | 0.60 | 6.8 |

**Supplementary Table 3:** Node and Edge Importance Distributions Across Training Stages.

| Metric | Model 1 (GAT Only) |  | Model 2 (GAT + BLC) |  |
| --- | --- | --- | --- | --- |
|  | Stage 1 | Stage 2 | Stage 1 | Stage 2 |
| **Node Importance** |  |  |  |  |
| Mean importance | $8.8 \times 10^{-4}$ | $1.1 \times 10^{-3}$ | $1.3 \times 10^{-3}$ | $3.6 \times 10^{-4}$ |
| Max importance | 0.066 | 0.077 | 0.20 | 0.031 |
| Excess kurtosis | 1080 | 1062 | 6393 | 330 |
| Gini coefficient | 0.48 | 0.45 | 0.47 | 0.42 |
| Bimodality<br>coefficient | 0.39 | 0.37 | 0.64 | 0.32 |
| **Edge Importance** |  |  |  |  |
| Mean importance | $9.5 \times 10^{-6}$ | $1.0 \times 10^{-5}$ | $2.6 \times 10^{-5}$ | $3.3 \times 10^{-6}$ |
| Max importance | $2.1 \times 10^{-3}$ | $3.7 \times 10^{-3}$ | $8.3 \times 10^{-3}$ | $1.8 \times 10^{-3}$ |
| Excess kurtosis | 865 | 1862 | 1264 | 5928 |
| Gini coefficient | 0.63 | 0.63 | 0.63 | 0.63 |

**Supplementary Table 4:** Ablation Analysis of Pathway Graph Structure on Model Performance. Table 4a: Effect of Edge and Node Removal on AD Classification Performance

| Experiment | Nodes | Edges | Edges/Node | $\Delta$ Edges | % $\Delta$ Stage 1 AUROC for AD classification | % $\Delta$ Stage 2 AUROC for AD classification |
| --- | --- | --- | --- | --- | --- | --- |
| Baseline (Pathway Graph) | 16, 352 | 938, 742 | 57 | N/A | 0.73 (0.70-0.76) | 0.78 (0.75-0.80) |
| Remove bottom 10 % of edges | 16, 352 | 844, 868 | 52 | -10% | -2.4% | -1.9% |
| Remove random 10 % of edges | 16, 352 | 844, 868 | 52 | -10% | -1.4% | -1.3% |
| Remove top 10 % of edges | 16, 352 | 845, 332 | 52 | -10% | -7.2% | -4.3% |
| Remove bottom 10 % of nodes | 14, 717 | 891, 423 | 63 | -5.0% | 0.0 % | 0.0 % |
| Remove random 10 % of nodes | 14, 717 | 766, 831 | 52 | -18% | 0.0 % | 0.0% |
| Remove top 10 % of nodes | 14, 717 | 675,715 | 46 | -28% | -4.1% | -4.1% |

**Table 4b:** Connectivity Characteristics of Top, Bottom, and Random Node Subsets.

| Group | N | Mean Degree | Median | 95% CI |
| --- | --- | --- | --- | --- |
| Top 10% Nodes | 1635 | 85 | 72 | [82, 88] |
| Bottom 10% Nodes | 1635 | 5.7 | 3.0 | [4.9, 6.7] |
| Random 10% | 1635 | 55 | 31 | [51, 58] |
| All Nodes | 16352 | 57 | 32 | [56.4, 58.5] |

**Table 4c:** Structural Impact of Edge Ablation on Pathway Graph.

| Graph | Edges | Components | Isolated Nodes (% $\Delta$) | Nodes w/ Degree = 1 (% $\Delta$) |
| --- | --- | --- | --- | --- |
| Original | 938, 742 | 3 | 0 | 8 (0.04%) |
| Remove Top 10% Edges | 844, 868 | 1920 | 1910 (11.7%) | 1510 (9.3%) |
| Remove Bottom 10% Edges | 844, 868 | 3 | 0 (0.0%) | 9 (0.05%) |
| Remove Random 10% Edges | 845, 332 | 22 | 19 (0.1%) | 434 (2.7%) |

**Supplementary Table 5:** 22 Concordant Genes across Fluid Intelligence and AD dimensions.

| Gene | Cohens_d_AD | Cohens_d_FI | FDR_AD | FDR_FI | z_AD | z_FI | abs_z_product | group |
| --- | --- | --- | --- | --- | --- | --- | --- | --- |
| APOE | 0.48 | 0.55 | 0 | 0 | 2.04 | 8 | 16.36 | case-driven |
| KDM2A | 0.42 | 0.47 | 0 | 0 | 1.38 | 7.18 | 9.94 | case-driven |
| ACTR3 | 0.16 | 0.21 | 0.02 | 0 | -1.53 | 4.67 | 7.14 | control-AD / case-FI |
| TBC1D26 | 0.16 | 0.15 | 0.02 | 0.04 | -1.57 | 4.09 | 6.42 | control-AD / case-FI |
| FHOD1 | 0.17 | 0.17 | 0.01 | 0.01 | -1.44 | 4.29 | 6.16 | control-AD / case-FI |
| MRPL17 | 0.41 | 0.17 | 0 | 0.01 | 1.29 | 4.3 | 5.53 | case-driven |
| SOCS5 | 0.42 | 0.13 | 0 | 0.04 | 1.37 | 3.96 | 5.42 | case-driven |
| OR51J1 | 0.2 | 0.19 | 0 | 0 | -1.13 | 4.51 | 5.1 | control-AD / case-FI |
| SCNN1D | 0.4 | 0.19 | 0 | 0 | 1.12 | 4.52 | 5.08 | case-driven |
| ZNF300 | 0.19 | 0.15 | 0 | 0.03 | -1.2 | 4.1 | 4.93 | control-AD / case-FI |
| SRF | -0.16 | -0.18 | 0.02 | 0.01 | -5.14 | 0.94 | 4.81 | control-driven |
| PPIA | -0.14 | -0.18 | 0.05 | 0.01 | -4.93 | 0.95 | 4.67 | control-driven |
| EGR1 | 0.38 | 0.22 | 0 | 0 | 0.89 | 4.81 | 4.28 | case-driven |
| MFSD8 | 0.39 | 0.15 | 0 | 0.03 | 1.02 | 4.12 | 4.2 | case-driven |
| TK2 | 0.21 | 0.15 | 0 | 0.03 | -0.98 | 4.09 | 4 | control-AD / case-FI |
| OR8U1 | 0.25 | 0.23 | 0 | 0 | -0.58 | 4.91 | 2.87 | control-AD / case-FI |
| DNAJC5G | 0.36 | 0.16 | 0 | 0.03 | 0.64 | 4.19 | 2.69 | case-driven |
| LINC00900 | 0.34 | 0.23 | 0 | 0 | 0.42 | 4.91 | 2.07 | case-driven |
| ATP6V0B | 0.34 | 0.19 | 0 | 0 | 0.43 | 4.53 | 1.96 | case-driven |
| C19orf25 | 0.26 | 0.15 | 0 | 0.02 | -0.4 | 4.13 | 1.66 | control-AD / case-FI |
| CCDC116 | 0.31 | 0.27 | 0 | 0 | 0.11 | 5.25 | 0.58 | case-driven |

**Supplementary Figure 1a:**
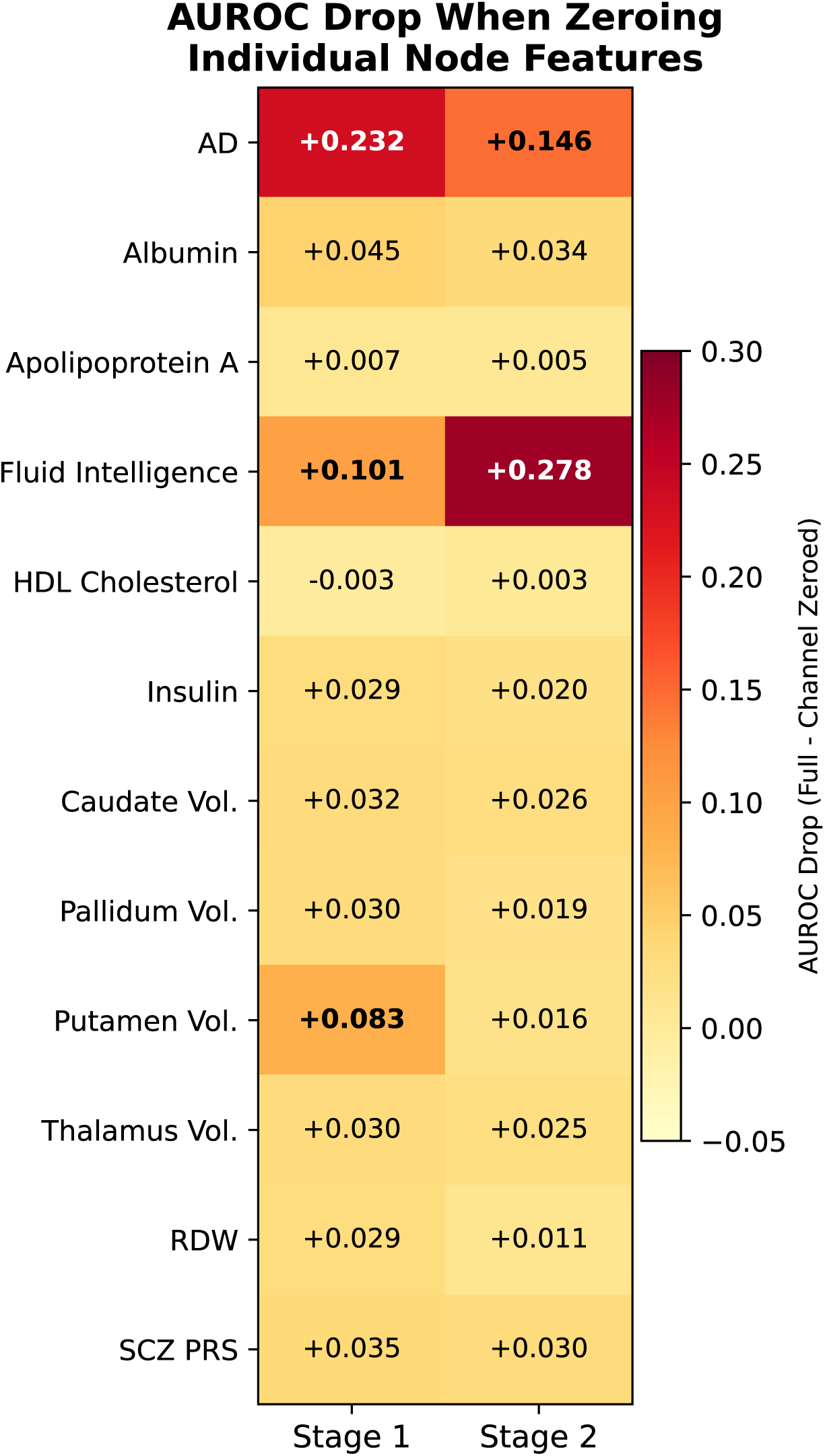
Impact of individual Node Feature Removal on AD Classification Accuracy

**Supplementary Figure 1b:**
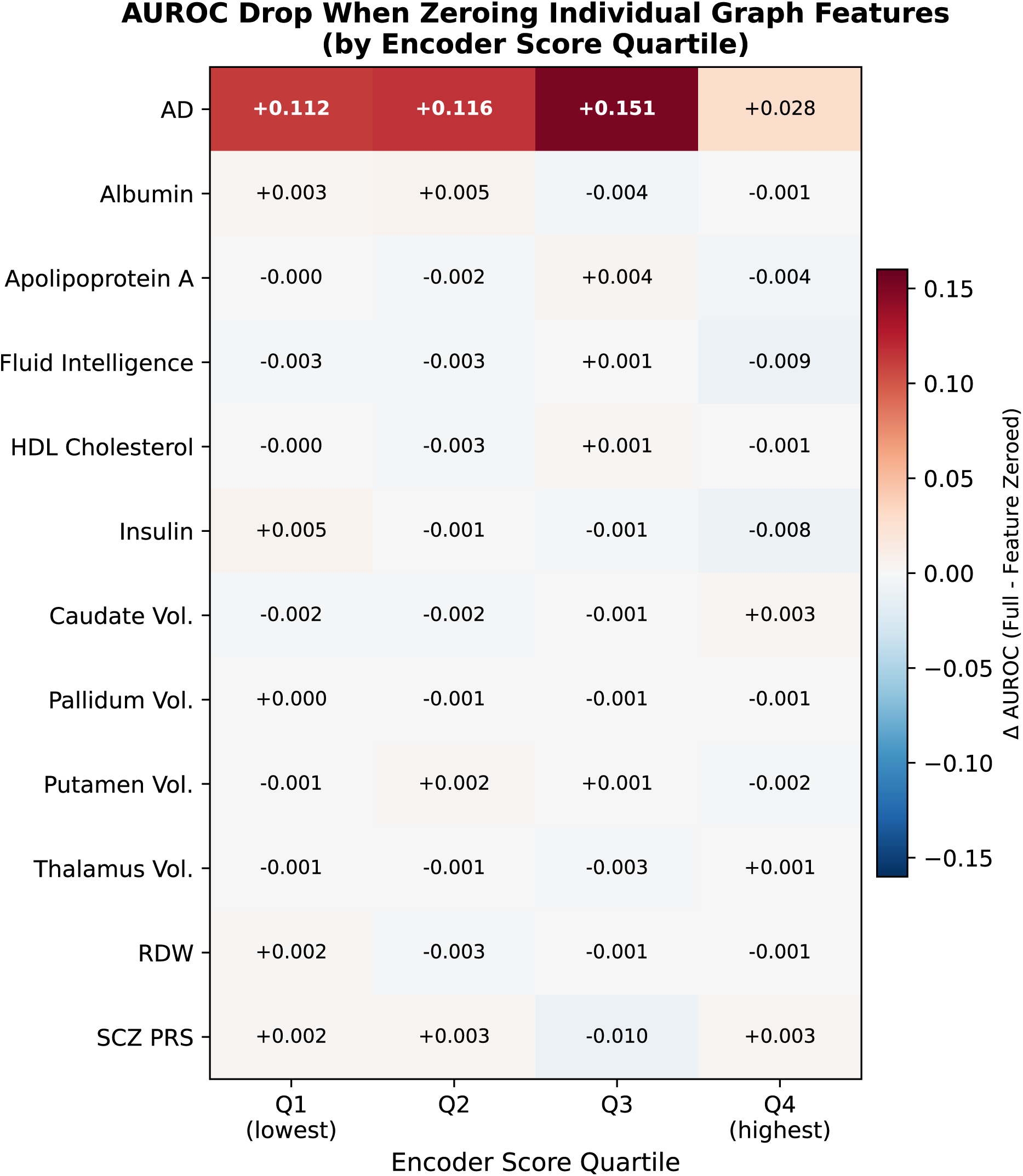
Impact of individual Graph Feature Removal on AD Classification Accuracy, stratified by Encoder Score Quartile

**Supplementary Figure 1c:**
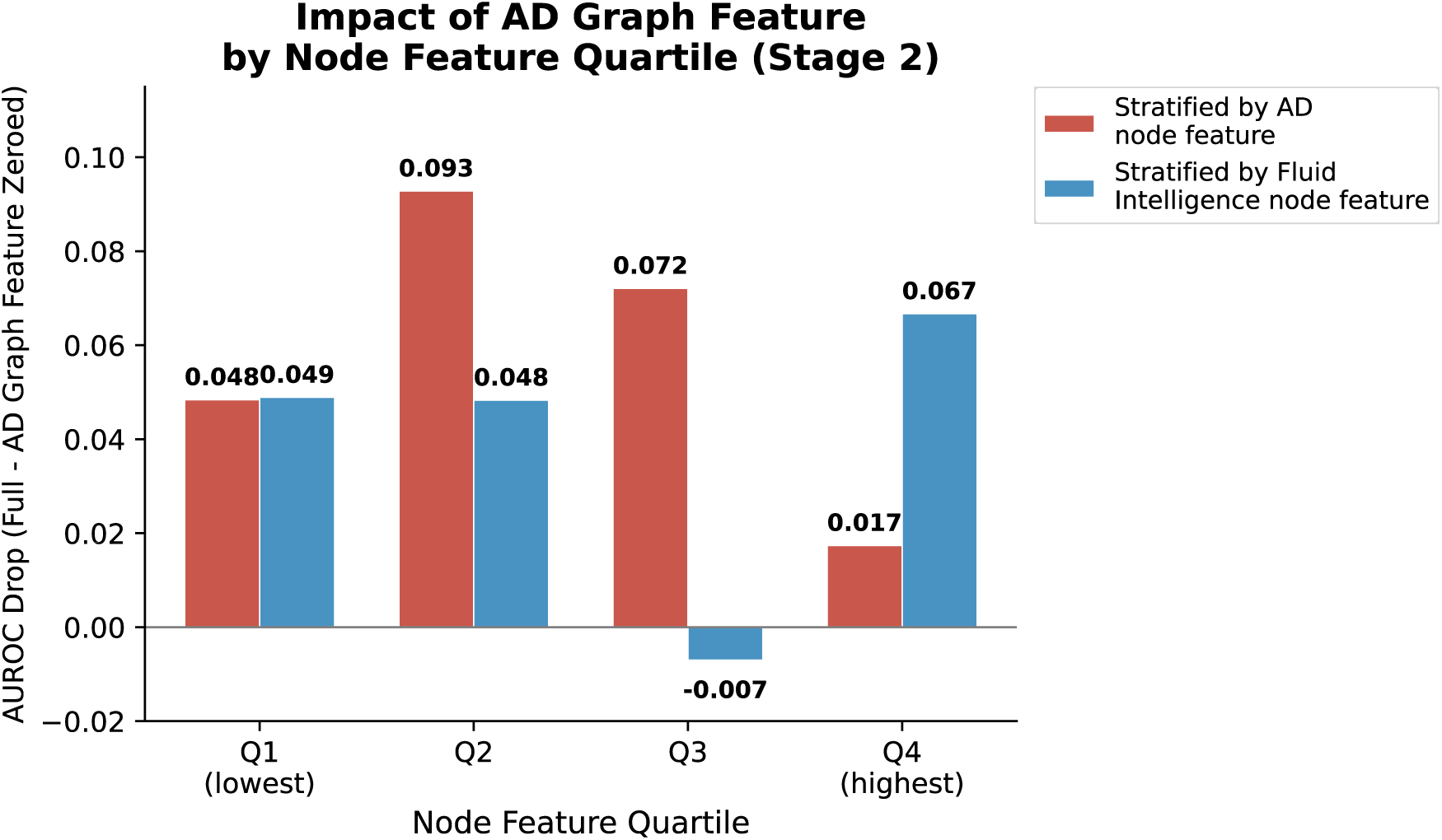
Impact of AD Graph Feature removal by Node Feature Quartile

**Supplementary Figure 2:**
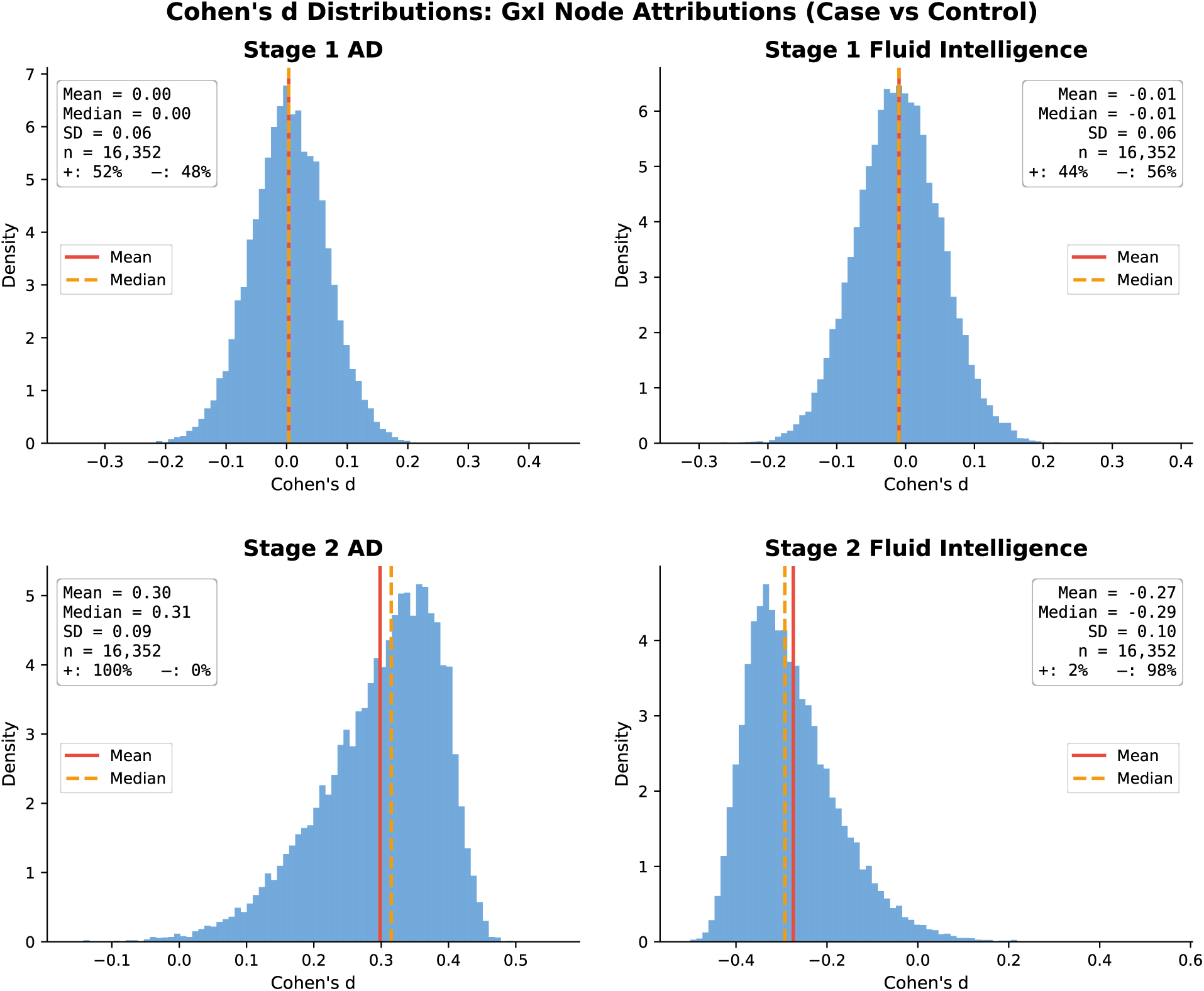
Cohen’s D Distributions for GxI Attributions for AD and Fluid Intelligence Node Features

## Notes

### Competing Interest Statement

Stephen Faraone received income, potential income, travel expenses continuing education support and/or research support from Aardvark, Aardwolf, AIMH, Akili, Atentiv, Axsome, Genomind, Ironshore, Johnson & Johnson/Kenvue, Kanjo, KemPharm/Corium, Noven, Otsuka, Sky Therapeutics, Sandoz, Supernus, Tris, and Vallon. With his institution, he has US patent US20130217707 A1 for the use of sodium-hydrogen exchange inhibitors in the treatment of ADHD. He also receives royalties from books published by Guilford Press: Straight Talk about Your Child's Mental Health, Oxford University Press: Schizophrenia: The Facts and Elsevier: ADHD: Non-Pharmacologic Interventions. He is Program Director of www.ADHDEvidence.org and www.ADHDinAdults.com. Dr. Faraone's research is supported by the European Union's Horizon 2020 research and innovation programme under grant agreement 965381; NIH/NIMH grants U01AR076092, R01MH116037, 1R01NS128535, R01MH131685, 1R01MH130899, U01MH135970, and Supernus Pharmaceuticals. His continuing medical education programs are supported by The Upstate Foundation, Corium Pharmaceuticals, Tris Pharmaceuticals and Supernus Pharmaceuticals. The other authors declare no competing interests.

### Funding

- Canadian Institutes of Health Research: 202110DFD-475191-95674

