## Supplementary File 4 for "Multi-Stage Graph Attention Networks for Interpretable Alzheimer’s Disease Classification from Genome-Wide Association Data"

Hyperparameters:

Stage 1:

{

"best_value": 0.7507300972938538,

"best_params": {

"repr_heads": 1,

"hidden_dim": 128,

"output_dim": 128,

"combined_scale": 0.7410543720018723,

"num_gat_layers": 2,

"dropout": 0.2,

"learning_rate": 0.0023655272133844387,

"weight_decay": 0.00036197376141578006,

"batch_size_s1": 32

},

"best_trial_number": 26,

"user_attrs": {

"best_auroc": 0.7507300979776536,

"class_weights": [

1.1313130855560303,

0.8960000276565552

],

}

Stage 2:

{

"best_value": 0.821711540222168,

"best_params": {

"lr_s2": 0.00043909805760996537,

"weight_decay_s2": 0.0008850928217491405,

"fusion_dim": 256,

"dropout_s2": 0.16777227554301052,

"freeze_gat": false,

"freeze_fc_gnn": true,

"batch_size_s2": 8

},

"best_trial_number": 9,

"user_attrs": {

"best_auroc": 0.8217115689381934,

"class_weights": [

1.1313130855560303,

0.8960000276565552

],

"gamma": 0.2,

"initial_batch_size": 8,

"successful_hyperparameters": {

"hidden_dim": 128,

"output_dim": 128,

"combined_dim": 96,

"num_gat_layers": 2,

"repr_heads": 1,

"learning_rate": 0.00043909805760996537,

"weight_decay": 0.0008850928217491405,

"fusion_dim": 256,

"dropout": 0.16777227554301052,

"freeze_gat": false,

"freeze_fc_gnn": true,

"initial_batch_size": 8,

"working_batch_size": 8

},

"working_batch_size": 8

}

}

Stage 3:

{

"best_value": 0.8223444819450378,

"best_params": {

"adv_lr": 0.001305127011606419,

"adv_grl_lambda": 1.0574753464833668,

"adv_weight_decay": 6.610628525240543e-05,

"adv_hidden_dim": 64,

"adv_dropout": 0.3812850400571666,

"lr": 0.0017774458871426598,

"weight_decay": 7.308788300773266e-05,

"batch_size_s3": 32

},

"best_trial_number": 11,

"user_attrs": {

"alpha": 0.2,

"best_auroc": 0.8223444866389579,

"best_mean_r2": -1.0,

"class_weights": [

1.1313130855560303,

0.8960000276565552

],

"gamma": 0.2,

"initial_batch_size": 32,

"successful_hyperparameters": {

"adv_lr": 0.001305127011606419,

"adv_grl_lambda": 1.0574753464833668,

"adv_weight_decay": 6.610628525240543e-05,

"adv_hidden_dim": 64,

"adv_dropout": 0.3812850400571666,

"learning_rate": 0.0017774458871426598,

"weight_decay": 7.308788300773266e-05,

"freeze_base": true,

"initial_batch_size": 32,

"working_batch_size": 32

},

"working_batch_size": 32

}

}
